# Peptidoglycan remodeling prevents antibiotic resistance during oxidative stress

**DOI:** 10.64898/2026.08.19.745765

**Authors:** Jonathan Sukadi Miala, Léanne Arcand-Carrier, Rosalie Lapointe, Claire Morin, Charles Sasseville, David Lalaouna, Eric Massé

## Abstract

The bacterial small RNA (sRNA) OxyS is expressed in *Escherichia coli* during oxidative stress. The sRNA OxyS enhances cell survival by controlling genes involved in the regulation of hydrogen peroxide (H_2_O_2_) and iron-sulfur (Fe-S) cluster formation. Here, we used the MS2 affinity purification coupled with RNA sequencing (MAPS) technique to identify new target mRNAs of the sRNA OxyS. Our analysis revealed a significant enrichment of *mepS* mRNA, which encodes a peptidoglycan endopeptidase that promotes cell growth. Our results confirm a previous report on the sRNA OxyS repressing the translation of *mepS*. We also found that an Δ*oxyS* background facilitates the emergence of mutations, conferring increased resistance to the last-resort antibiotics polymyxin B and E (colistin), but only in the presence of the target *mepS* gene. This suggests that the translation repression of *mepS* by OxyS could prevent mutations in bacterial DNA during H_2_O_2_-induced oxidative stress. Moreover, we show that adding the antioxidant thiourea or sequestering iron in the Δ*oxyS* background effectively reduces the emergence of resistance against both polymyxin B and colistin. These results suggest that reactive oxygen species (ROS), in conjunction with intracellular iron, play a key role in driving the emergence of antibiotic resistance. Overall, our work underlines a mechanism of antimicrobial emergence implicating oxidative stress, intracellular Fe, and cell wall remodeling in *E. coli*.

**IMPORTANCE:** This study uncovers an underexplored link between peptidoglycan remodeling and oxidative stress responses during exposure to antibiotics. By elucidating how MepS and the sRNA OxyS interact in the presence of polymyxins and oxidative stress, our study suggests that MepS may exert an anti-mutator function. The repression of *mepS* translation by OxyS seems to limit the emergence of antibiotic resistance driven by DNA mutations. Together, these findings suggest cell wall remodeling and oxidative stress response pathways as promising targets to enhance antibiotic efficacy and limit the emergence of resistance.

## INTRODUCTION

To cope with constantly changing environmental conditions, bacteria rely on small regulatory RNAs (sRNAs), which rapidly adjust gene expression in response to stress [1] [2]. sRNAs are regulatory molecules of approximately 100 to 200 nucleotides in length that are expressed under challenging environmental conditions, including virulence conditions, and can help bacterial cells invade the host [1] [3]. These molecules can modify the expression of certain genes by pairing with their target mRNAs at various sites, such as the ribosome binding site (RBS), 5’ or 3’ UTRs, as well as within the coding sequence (CDS) [1]. For stable pairing with a target mRNA, sRNAs may require binding of chaperone proteins, such as Hfq or ProQ [4] [5] [6].

One of the primary challenges that bacteria encounter is oxidative stress, which originates from the accumulation of reactive oxygen species (ROS), such as hydroxyl (^●^OH) and superoxide (O_2_•-) radicals [7]. These ROS stem from the many metabolic activities of bacterial cells, for instance cellular respiration [7] [8]. The accumulation of ROS can also result from the external environment. For example, H_2_S production by sulfur reducing bacteria can react with O_2_ in the intestinal epithelium to produce H_2_O_2_ in the intestinal space [7]. Moreover, during the process of phagocytosis by macrophages, the engulfed bacteria are exposed to high concentration of H_2_O_2_, which induces oxidative stress [7]. While H_2_O_2_ is mildly toxic, it also acts as a precursor to more deleterious species via the Fenton reaction, where intracellular iron (Fe) reacts with H_2_O_2_ to produce hydroxyl radicals (^●^OH). The highly toxic ^●^OH can damage proteins and lipids as well as induce mutations in RNA and DNA [9] [10]. Moreover, it has been suggested that antibiotic treatments can increase the level of ROS in bacteria [11] [12]. The Oxidative stress response was also shown to aid bacterial survival and adaptation to antibiotics and may influence the emergence of *de novo* antibiotic resistance [13] [14].

Antibiotic resistance is an increasingly serious concern, undermining the effectiveness of treatments for bacterial infections [15]. The rise in bacterial resistance can lead to complications, including prolonged hospitalizations, increased healthcare costs, treatment failures, and preventable deaths [15]. Although oxidative stress can promote adaptation to antibiotics, bacterial cells limit the level of endogenous ROS through dedicated stress sensors such as the transcription factor OxyR in *E. coli* [16] [17]. OxyR orchestrates the oxidative stress response by inducing the expression of genes such as *oxyS*, which encodes an sRNA involved in mitigating intracellular ROS levels [16] [17].

OxyS is a 117-nt long sRNA that plays a role in gene regulation during oxidative stress and exerts an anti-mutator effect in *E. coli* [18] [19] [20]. The sRNA OxyS helps bacteria repair DNA damage, thereby limiting the occurrence of mutations [21]. To achieve this, OxyS represses the transcriptional factor *nusG*, which leads to cell division arrest, allowing the bacteria to repair DNA damage before resuming division [21]. Additionally, OxyS regulates the cellular response to intracellular H_2_O_2_ generation [22]. The expression of OxyS can also directly increase the translation of *iscR* transcription regulator of the *iscRSUA* operon, which is involved in the biosynthesis of Fe-S clusters [23]. Recently, another group used RIL-seq (RNA interaction by ligation and sequencing) to characterize the interaction of OxyS with the *mepS* mRNA near the ribosome binding site (RBS), which repressed the translation of the mRNA *mepS* during oxidative stress [33].

The *mepS* mRNA encodes the lipoprotein MepS, encoding a D-D peptidoglycan endopeptidase that contributes to the growth of the peptidoglycan cell wall in *E. coli*. The peptidoglycan cell wall, together with the outer membrane and the inner membrane, form the bacterial cell envelope. The bacterial cell wall is a dynamic structure, constantly adapting to the environment, and composed of peptidoglycan layers [24] [25] [26] [27]. The MepS protein facilitates the bacterial cell elongation during growth by cleaving peptide crosslinks in the mature peptidoglycan cell wall. This cleavage generates space for the incorporation of new peptidoglycan, enabling cell wall expansion and proper bacterial elongation [28] [29]. The bacterial peptidoglycan cell wall surrounds most Gram-positive and Gram-negative bacteria, giving them shape and protection against lysis from osmotic pressure. Because of its importance, the peptidoglycan cell wall is a preferred target in various antibacterial therapeutic interventions [30]. Indeed, the synthesis of peptidoglycans is often targeted by antibiotics, such as beta-lactams, which inhibit the final steps of peptidoglycan synthesis by acylating the transpeptidases responsible for cross-linking peptides [25].

To investigate the interactome of the sRNA OxyS and find additional mRNA targets, we performed a MAPS (MS2-affinity purification coupled with RNA sequencing) on a MS2-OxyS construct [31] [32]. We found a significant enrichment of the mRNA *mepS* associated with MS2-OxyS in *E. coli*. Here, we used experiments such as probing, gene fusions, northern blots, and western blots to characterize the interaction between OxyS and the mRNA *mepS.* We observed an interaction of the sRNA OxyS on *mepS* mRNA near the ribosome binding site, which repressed the translation of the mRNA *mepS* during oxidative stress, confirming a recent report using RIL-seq [33]. We observed that decreased expression of MepS protein significantly disrupts bacterial cell envelope integrity as indicated by chlorophenol red-β-D-galactopyranoside (CPRG) assays, a chemical dye used to detect cell damage [58]. We also provide evidence that the absence of the sRNA OxyS promotes resistance against polymyxin B and E (colistin), two major antibiotics used as last resort drugs in clinics. Our study may suggest that a potential interaction between OxyS and mRNA *mepS* during oxidative stress could potentially result in an anti-mutator effect as the absence of MepS prevents DNA mutations and the emergence of polymyxin resistance. Additional data indicated that the antioxidant thiourea or iron chelator 2.2’-dipyridyl effectively reduced the emergence of resistance against both polymyxin B and colistin in the Δ*oxyS* background. This suggests that ROS, probably generated through intracellular iron, play a role in driving the emergence of antibiotic resistance. Our work underlines a previously unsuspected role for the MepS endopeptidase, by remodeling cell wall integrity during oxidative stress, to prevent mutations and the emergence of antimicrobial resistance.

## RESULTS

### Oxidative stress and the sRNA OxyS modulate the level of MepS protein

We first sought to find new RNA targets of the sRNA OxyS by performing a MAPS experiment on MS2-OxyS construct. After pull-down of MS2-OxyS and sequencing of co-purified RNAs, we identified *mepS* as a promising binding partner of the sRNA OxyS (Fig.1A) among a variety of potential target mRNAs (GEO number: to be determined). The bedgraph from MS2-OxyS MAPS, showing RNA-seq coverage, indicates strong enrichment of the *mepS* transcript in the vicinity of the ribosome binding site (RBS) (Fig 1A). This suggests that the sRNA OxyS could potentially bind near the translation initiation of the *mepS* mRNA and regulate ribosome binding. To confirm this, we performed an in vitro lead acetate probing assay and found that the sRNA OxyS binds to *mepS* partly on the Shine-Dalgarno sequence (Fig. S1). To investigate this pairing, we made nucleotides changes at the interaction site (seed binding) of the sRNA OxyS and mRNA *mepS+192* LacZ (Fig. S2D) to create the *mepSmut* construct and disrupt the interaction. We then measured the beta-galactosidase activity of this MepSmut+192-LacZ translational fusion to monitor the interaction between OxyS and *mepS* in cells. Our results show that WT OxyS represses WT MepS +192-LacZ and that OxySmut represses MepSmut+192-LacZ (Fig. S3). This suggests that OxyS could repress MepS+192-LacZ translation and that the interaction occurs at the site suggested in Fig 1A. Our results corroborate a previous report characterizing the interaction between OxyS and *mepS* mRNA [33].

**Figure 1.**
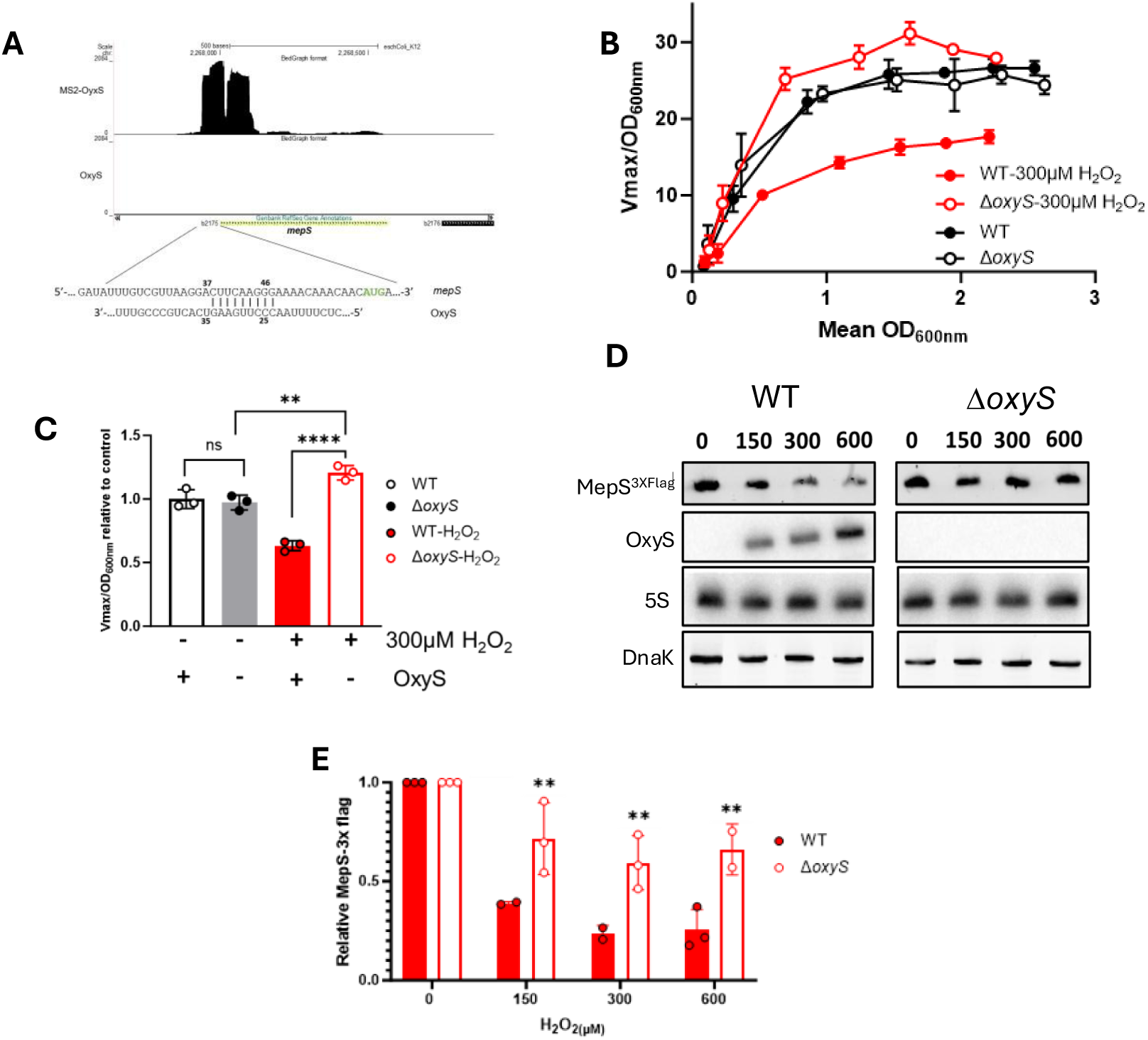
H_2_O_2_ modulates the level of MepS protein through the expression of OxyS. **(A)** Bedgraph of MS2-OxyS construct and OxyS control from the MAPS assay. The enrichment ratio of MS2-OxyS/OxyS is 1638. **(B)** β-galactosidase assays of translational MepS+192-LacZ construct from WT or Δ*oxyS* backgrounds in LB medium. Expression of the sRNA OxyS was induced via 300 μM of H_2_O_2_ every 15 minutes after 1 hour of growth (n=3). **(C)** Relative activity of β-galactosidase assays from (B) translational MepS+192-LacZ construct from WT and Δ*oxyS* backgrounds. Expression of the sRNA OxyS was induced every 15 minutes via 300 μM of H_2_O_2_ after 1 hour of growth. Samples were taken at OD_600nm_=1.5 for relative activity (n=3). Statistical analysis were performed using ordinary one-way ANOVA : Tukey s multiple comparisons test: ns not significant p-value: 0.9351, * (p-value < 0.05), ** (p-value < 0.01), *** (p-value < 0.001), **** (p-value < 0.0001). (**D)** Western blot of MepS-3xFlag construct from WT or Δ*oxyS* backgrounds grown in LB medium. Expression of OxyS was induced with 150, 300, and 600 μM of H_2_O_2_ at OD_600nm_=0.5. Samples were taken after induction of OxyS. DnaK and 5S rRNA were used as loading controls for MepS^3XFlag^ tagged protein and sRNA OxyS, respectively (n=3). **(E)** Relative expression of MepS^3XFlag^ following induction of the sRNA OxyS with H_2_O_2_ (150, 300, and 600 M) at OD_600nm_=0.5.

Following this, we studied the expression of the protein MepS in the presence of oxidative stress. To address this, we used our MepS+192-LacZ translational fusion in WT and Δ*oxyS* backgrounds in the presence or in the absence of H_2_O_2_ (Fig. 1B and C). In the absence of H_2_O_2_, the expression of MepS+192-LacZ fusion in the WT strain and the mutant Δ*oxyS* was similar (Fig. 1B) (see Fig. S2A and B for endogenous expression of OxyS). However, when we added 300 µM of H₂O₂ to the culture every 15 min, the activity of MepS+192-LacZ fusion was significantly lower (57% lower) in the WT strain compared to the Δ*oxyS* mutant (Fig. 1B-C, see supp Fig. 2C for level of sRNA OxyS after induction with H_2_O_2_). Results show no significant difference in the MepS+192-LacZ fusion activity between the WT strain and the Δ*oxyS* mutant in the absence of H_2_O_2_ (Fig. 1C). Following this, we monitored the effect of the sRNA OxyS on the MepS^3xFlag^ tagged protein. We performed a western blot on MepS^3xFlag^ in parallel with a northern blot to monitor the level of the sRNA OxyS after induction with H_2_O_2_. We observed a dose-dependent increase in sRNA OxyS expression level following induction with H_2_O_2_ in the WT background (Fig. 1D). In contrast, the level of tagged protein MepS^3xFlag^ decreases in a WT strain compared to the Δ*oxyS* mutant as shown by western blot (Fig. 1D). The decrease in MepS^3xFlag^ expression correlates with the increased level of cellular OxyS. We then quantified the cellular level of the protein MepS^3xFlag^ and noticed a significant decrease in WT strain compared to the mutant Δ*oxyS* (Fig. 1E).

**Figure 2.**
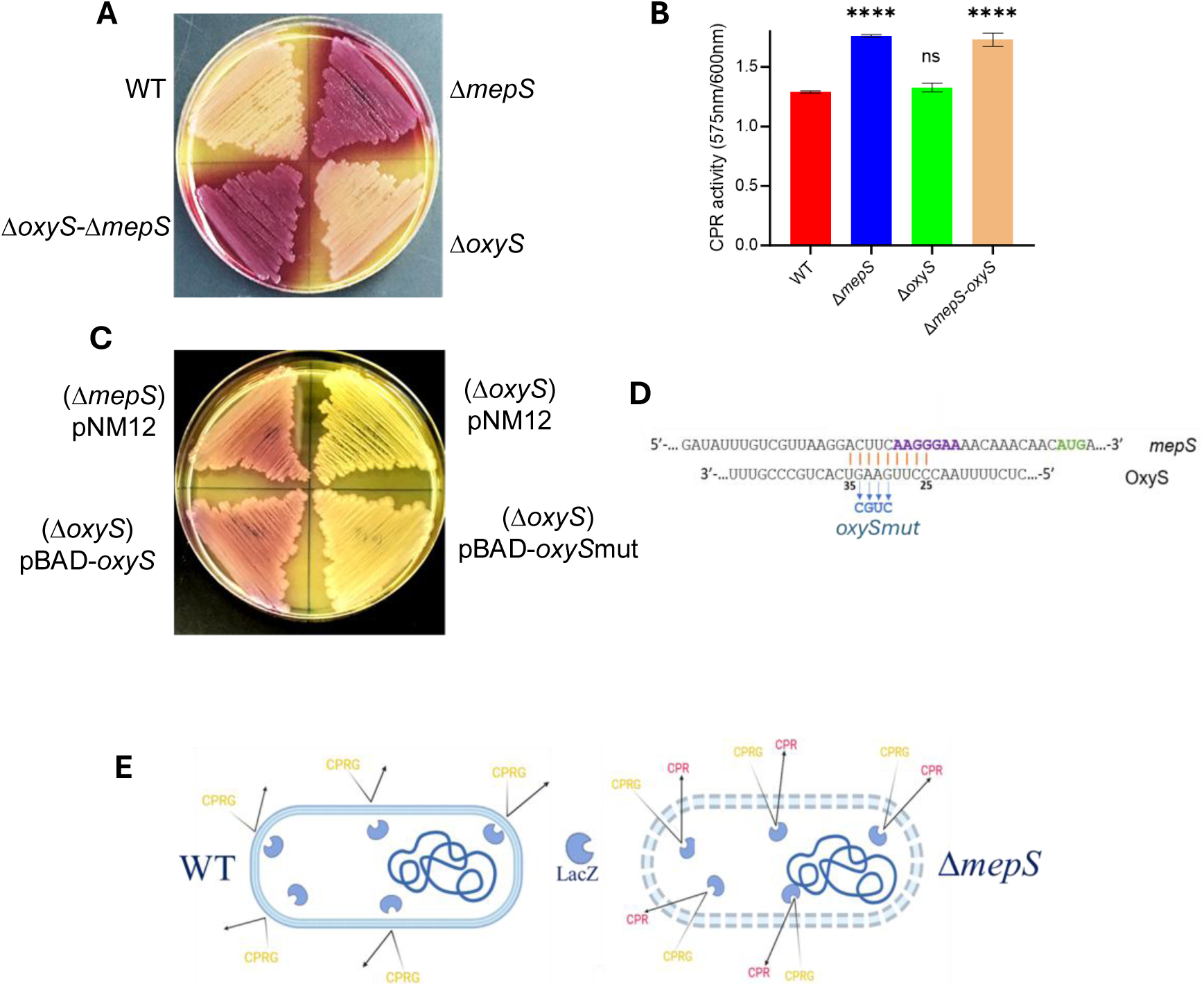
The sRNA OxyS-*mepS* interaction affects the membrane integrity of *E. coli.* **(A)** CPRG assay for WT, Δ*mepS*, Δ*oxyS,* and Δ*mepS-*Δ*oxyS* cells grown on solid LB0N (LB without NaCl) with 200µg/mL CPRG. **(B)** CPRG assay for WT, Δ*mepS*, Δ*oxyS,* and Δ*mepS-*Δ*oxyS* cells grown in liquid LB0N with 30µg/mL CPRG. Samples were taken after15 min incubation (CPR activity = 575nm/600nm) (n=3). Statistical analysis were performed using One-way ANOVA : Dunnett’s multiple comparisons test: ns not significant : 0.3369, **** (p-value < 0.0001). **(C)** CPRG assay in LB0N (LB without NaCl) with 200µg/mL CPRG and 0.1% of arabinose in strains carrying pNM12 or pBAD-OxyS (Δ*mepS* or Δ*oxyS* backgrounds). **(D)** Nucleotide modification site for mutant OxySmut sRNA. (**E)** Schematic representation of the CRPG assay adapted from (58)

### The sRNA OxyS pairing with the mRNA *mepS* affects membrane integrity of *E. coli*

The sRNA OxyS targets *mepS* mRNA, which encodes an enzyme critical for cell wall remodeling and for maintaining the permeability of the bacterial envelope [28][50]. This regulation suggests that the interaction between OxyS and mRNA *mepS* could influence the cellular permeability under conditions of OxyS expression. To address this, we used chlorophenol red-β-D-galactopyranoside (CPRG) product, a B-galactosidase enzyme substrate which typically cannot penetrate the envelope of gram-negative bacterial cells [58]. However, when envelope integrity is compromised, cells become more permeable to CPRG, allowing the substrate to enter the cytoplasm and react with the endogenous LacZ enzyme to produce the red compound chlorophenol red (CPR). In addition, the CPRG assay also detects the release of LacZ from lysed cells, allowing to measure both envelope permeability to CPRG and the propensity of cells to undergo lysis. (Fig. 2E). For these experiments we used LB medium without NaCl (LB0N) because the mutant Δ*mepS* was previously reported to be sensitive to osmotic stress [50]. First, on solid LB0N medium, we observed that both the Δ*mepS* mutant and the double Δ*mepS-*Δ*oxyS* mutant produced red colonies, suggesting that CPRG was able to penetrate into the cytoplasm due to increased envelope permeability and that a fraction of the bacterial population may also have undergone lysis, releasing endogenous LacZ. These findings are consistent with impaired envelope integrity (Fig. 2A). In contrast, the WT strain and the Δ*oxyS* mutant produced yellow colonies indicating that CPRG could not reach the cytoplasm and LacZ enzyme. These phenotypes suggest that the membrane integrity for these strains (WT and Δ*oxyS* mutant) was intact and the absence of the sRNA OxyS does not have an impact on the membrane integrity comparatively to the Δ*mepS* background (Fig. 2A). We next quantified the CPR level of our strains using a spectrophotometer (575 nm / 600 nm) as performed previously [63]. The mutant Δ*mepS* and double mutant Δ*mepS-*Δ*oxyS* showed the higher level of CPR compared to the WT strain and *ΔoxyS* mutant (Fig. 2B). Finally, to study the effect of the interaction between OxyS and its target *mepS* mRNA on the membrane integrity we induced the expression of the OxyS sRNA from a pBAD-OxyS plasmid in a Δ*oxyS* background strain (Fig. 2C). The overexpression of OxyS generated noticeably red colonies similar to the control strain Δ*mepS* with pNM12 (empty plasmid control). This suggests that the interaction between OxyS and *mepS* mRNA could potentially affect membrane integrity. Also, to investigate if this phenotype was specific to the sRNA OxyS-*mepS* interaction, we induced the expression of the mutated OxySmut construct from a pBAD plasmid (Fig. 2C and 2D). Fig. 2C shows that cells expressing OxySmut (Fig. 2D) produced yellow colonies, like the control Δ*oxyS* strain with pNM12, which suggests that the previous phenotype was due to the specific interaction between OxyS and the target mRNA *mepS* (Fig. 2C). Overall, these results suggest that the specific interaction between OxyS and *mepS* reduces MepS protein expression, which may compromise cell envelope integrity and increase envelope permeability.

### The OxyS–*mepS* interaction potentially drives polymyxin B and colistin resistance

Given that the Δ*mepS* mutant exhibited compromised cell wall integrity, we next assessed the susceptibility of our mutant strains to cell envelope-targeting antibiotics. These antibiotics, such as mecillinam, vancomycin, polymyxin B, and colistin disrupt either the peptidoglycan cell wall or the LPS-containing outer membrane. We first exposed the WT, Δ*mepS*, Δ*oxyS*, and the Δ*mepS-*Δ*oxyS* double mutant strains to increase concentrations of mecillinam. At 0.125 µg/mL of mecillinam, we observed that the Δ*mepS* mutant, and the Δ*mepS-*Δ*oxyS* double mutant exhibited complete growth inhibition, compared to the WT and Δ*oxyS* strains where we observed growth (Fig. 3A). However, we observed an absence of growth at a concentration of 0.25 µg/mL of mecillinam for the WT and Δ*oxyS* strains (Fig. 3A). These results suggest that absence of *mepS* could make strains more sensitive than a WT in the presence of mecillinam antibiotic, which corroborated with a previous study [51]. We also used the cell wall inhibitor vancomycin and found that the mutant Δ*mepS* was more sensitive than WT or Δ*oxyS* at concentrations of 100 and 150 µg/mL (Fig. S5C). Moreover, to determine whether the compromised envelope could confer broader sensitivity to unrelated antibiotic classes, we included rifampicin, which inhibits RNA polymerase and does not target the cell envelope. We noticed that at concentration of 25 µg/mL of rifampicin, the Δ*mepS* mutant and the Δ*mepS-*Δ*oxyS* double mutant were more sensitive to rifampicin than the WT and Δ*oxyS* strains. In contrast, at 100 µg/mL rifampicin, no growth was observed for any of the strains (Fig. S5A). These results suggest that the Δ*mepS* mutant may exhibit an increased sensitivity to diverse antibiotic classes beyond those specifically targeting the cell wall.

**Figure 3.**
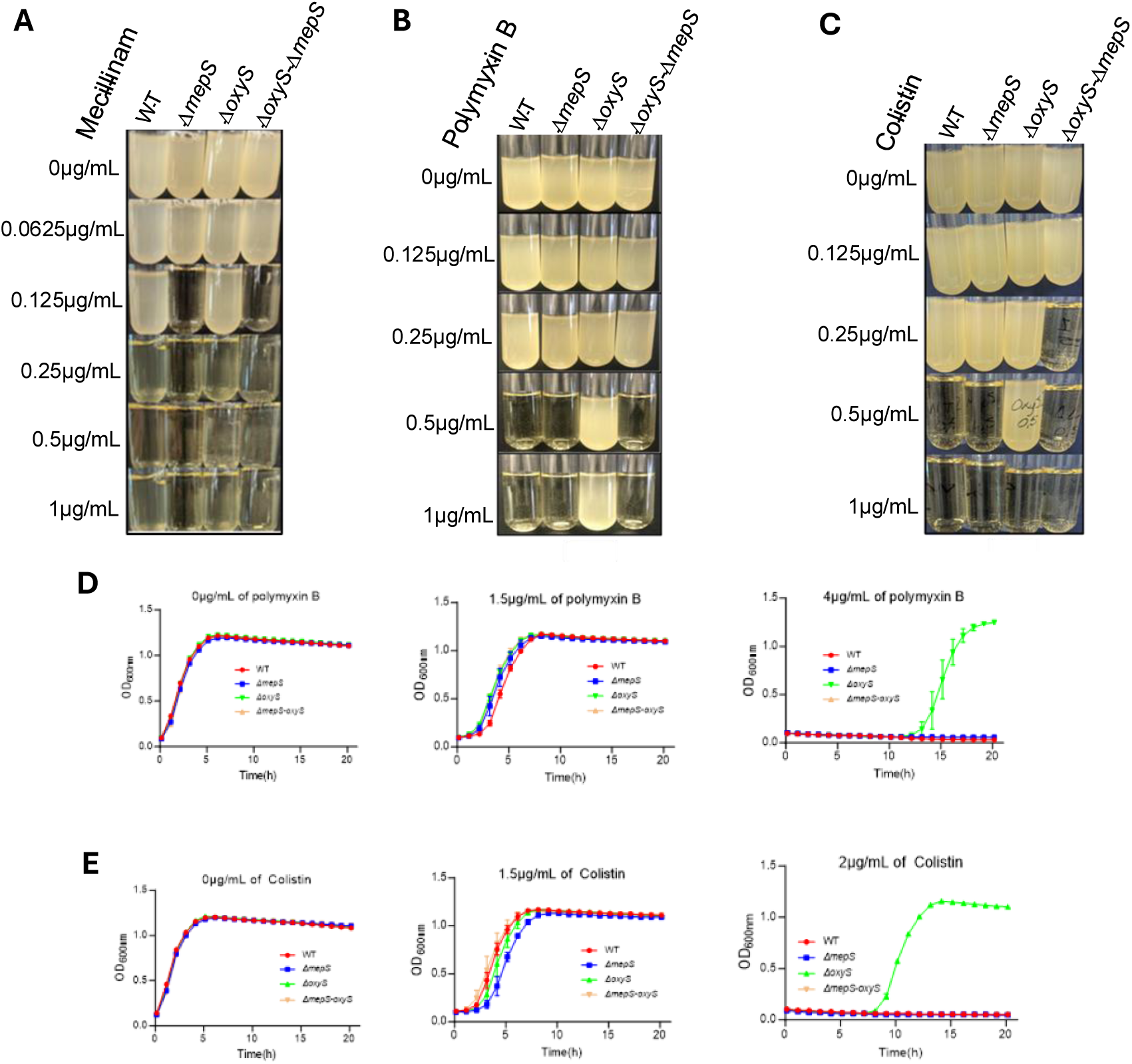
The sRNA OxyS is involved in resistance against polymyxin B and colistin antibiotics. **(A)** Growth assay at 37°C in LB medium of WT, Δ*mepS*, Δ*oxyS,* and Δ*mepS-*Δ*oxyS* cells in presence of the antibiotic mecillinam at different concentrations (0, 0.0625, 0.125, 0.25, 0.5, and 1µg/mL) (n=3). **(B)** Growth assay of WT, Δ*mepS*, Δ*oxyS,* and Δ*mepS-*Δ*oxyS* cells at 37°C in LB medium in the presence of the antibiotic polymyxin B at different concentrations (0, 0.125, 0.25, 0.5, and 1µg/mL) (n=3). **(C)** Growth assay of WT, Δ*mepS*, Δ*oxyS,* and Δ*mepS-*Δ*oxyS* cells at 37°C in LB medium in presence of the antibiotic colistin at different concentrations (0, 0.125, 0.25, 0.5, and 1µg/mL) (n=3). **(D)** Growth curve assay of WT, Δ*mepS*, Δ*oxyS,* and Δ*mepS-*Δ*oxyS* cells in 96-well microplates at 37°C in LB medium in the presence of polymyxin B at different concentrations (0, 1.5, and 4 µg/mL). All growth curves start with samples at OD_600nm_=0.1 (n=3). **(E)** Growth curve assay of WT, Δ*mepS*, Δ*oxyS,* and Δ*mepS-*Δ*oxyS* cells in 96-well microplates at 37°C in LB medium in the presence of colistin at different concentrations (0, 1.5, and 2 µg/mL) (n=3).

We further assessed the impact of the Δ*mepS* mutant on antibiotic susceptibility and tested additional antibiotics that act on the bacterial cell envelope, namely polymyxin B and colistin. Polymyxin B and colistin are cationic molecules that bind the bacterial lipopolysaccharide (LPS) at the sub-unit level called Lipid A, thereby disrupting the bacterial membranes and inducing cell lysis [34]. We used a WT strain, mutant Δ*mepS,* mutant Δ*oxyS*, and a double mutant Δ*mepS-*Δ*oxyS* in the presence of different concentrations of polymyxin B and we noticed that all strains but the mutant Δ*oxyS* stopped growing at 0.5 µg/mL of polymyxin B (Fig. 3B). The Δ*oxyS* mutant could grow at 0.5 µg/mL and even in the presence of 1 µg/mL of polymyxin B (Fig. 3B). We then asked whether the phenotype observed in the Δ*oxyS* mutant was due to its interaction with the target mRNA *mepS*. To achieve this, we mutated the interaction site of the sRNA OxyS (from CUUC to GCAG), thereby creating OxySmut (Fig. S4B). We observed that the strain with a mutated interaction site in OxyS grew better than the WT strain. Indeed, the WT strain stopped growing at 0.5 µg/mL of polymyxin B, but the strain with an OxyS mutated in the interaction site continues to growth until 1 µg/mL of polymyxin B (Fig. S4A). We also monitored the effect of colistin in our strains (WT, Δ*mepS*, Δ*oxyS*, and Δ*mepS-*Δ*oxyS*). Again, the Δ*oxyS* mutant displayed a higher tolerance to colistin, maintaining growth up to 0.5 µg/mL (Fig 3C), while the remaining strains exhibited lower resistance thresholds: the WT and Δ*mepS* strains ceased growth at 0.5 µg/mL, while the Δ*mepS-*Δ*oxyS* double mutant stopped at 0.25 µg/mL.

Following this, we performed growth curve assays with our strains (WT, Δ*mepS*, Δ*oxyS*, and Δ*mepS-*Δ*oxyS*) in the presence of polymyxin B at various concentrations. We observed a slight negative effect on growth starting at 1.5 µg/mL of polymyxin B on all our strains (Fig. 3D). Although no growth was observed during the first ∼10 hrs of incubation at 4 µg/mL of polymyxin B, the Δ*oxyS* mutant strain exhibited robust growth after 12 hours of incubation at this concentration (Fig. 3D). Next, we aimed to monitor the effect of antibiotic colistin on our mutant strains (Fig. 3E). From 0 to 1.5 µg/mL of colistin, we observed again a reduced growth on all strains (Fig. 3E). While there was no noticeable growth at 2µg/mL of colistin from 0 hour to 8 hours, strong growth became apparent for the Δ*oxyS* mutant after 8 hours (Fig. 3E). Overall, these results suggest that the mutant Δ*oxyS* can develop resistance to polymyxin B and colistin through time compared to WT, Δ*mepS*, and Δ*oxyS*-Δ*mepS* backgrounds. This also suggests that the *mepS* gene seems to be involved in the mechanism of resistance from the Δ*oxyS* mutant (Fig. 3D and E, compare Δ*oxyS* with Δ*oxyS*-Δ*mepS* backgrounds).

### Emergence of resistance to polymyxin B and colistin in Δ*oxyS* mutant may depend on ROS

To explain the results in Fig. 3, we hypothesized that the Δ*oxyS* background leads to increased levels of endogenous oxidative stress and the accumulation of ROS [13] [14] [18] [21]. The presence of ROS, such as ^●^O_2_^-^ and ^●^OH, was previously shown to promote mutation rate, thereby facilitating the emergence of antibiotic resistance [13] [14]. To investigate this hypothesis, we used thiourea, which is an antioxidant and scavenger of ^●^O_2_^-^and ^●^OH (Fig. 4C), and performed an experiment similar to Fig. 3B-C in the presence of polymyxin B and colistin. We first incubated a WT strain and Δ*oxyS* mutant in the presence of polymyxin B or colistin at 0.4 µg/mL (Fig. 4A). Only the Δ*oxyS* mutant was able to growth in that condition compared to the WT strain, which did not grow in presence of antibiotics. However, when we added 50mM of the antioxidant thiourea to the cultures with polymyxin B or colistin, neither the WT nor the Δ*oxyS* were able to grow (Fig. 4A). Next, we used the iron chelator 2.2’-dipyridyl, which sequesters intracellular Fe^2+^ and thereby prevents ^●^OH production from the Fenton reaction (Fig. 4C). We incubated both WT and Δ*oxyS* mutant strains in the presence of polymyxin B or colistin (0.4 µg/mL) and in the presence or in the absence of 2.2’-dipyridyl at 0.25 mM (Fig. 4B). When we added the 2.2’-dipyridyl to the culture with the antibiotic, the Δ*oxyS* mutant was unable to grow in these conditions just like the WT strain, suggesting that polymyxin B and colistin are killing cells (Fig. 4B). Moreover, we quantified the level of ROS with HPF fluorescein dye, which specifically detects ^●^OH [11]. The HPF assays indicated that the mutant Δ*oxyS* had significantly more fluorescence signal (∼18%) than the WT strain during treatment with 1 µg/mL of polymyxin B (Fig. 4D) and 17 % more with colistin (Fig. 4E). We also observed that the presence of thiourea (50mM) decreased the HPF fluorescein signal as compared to (Fig. 4D and E). These results may indicate an increased level of ROS in the Δ*oxyS* mutant background and suggest that ROS could play a role in the development of resistance to polymyxin B and colistin in a Δ*oxyS* mutant.

**Figure 4.**
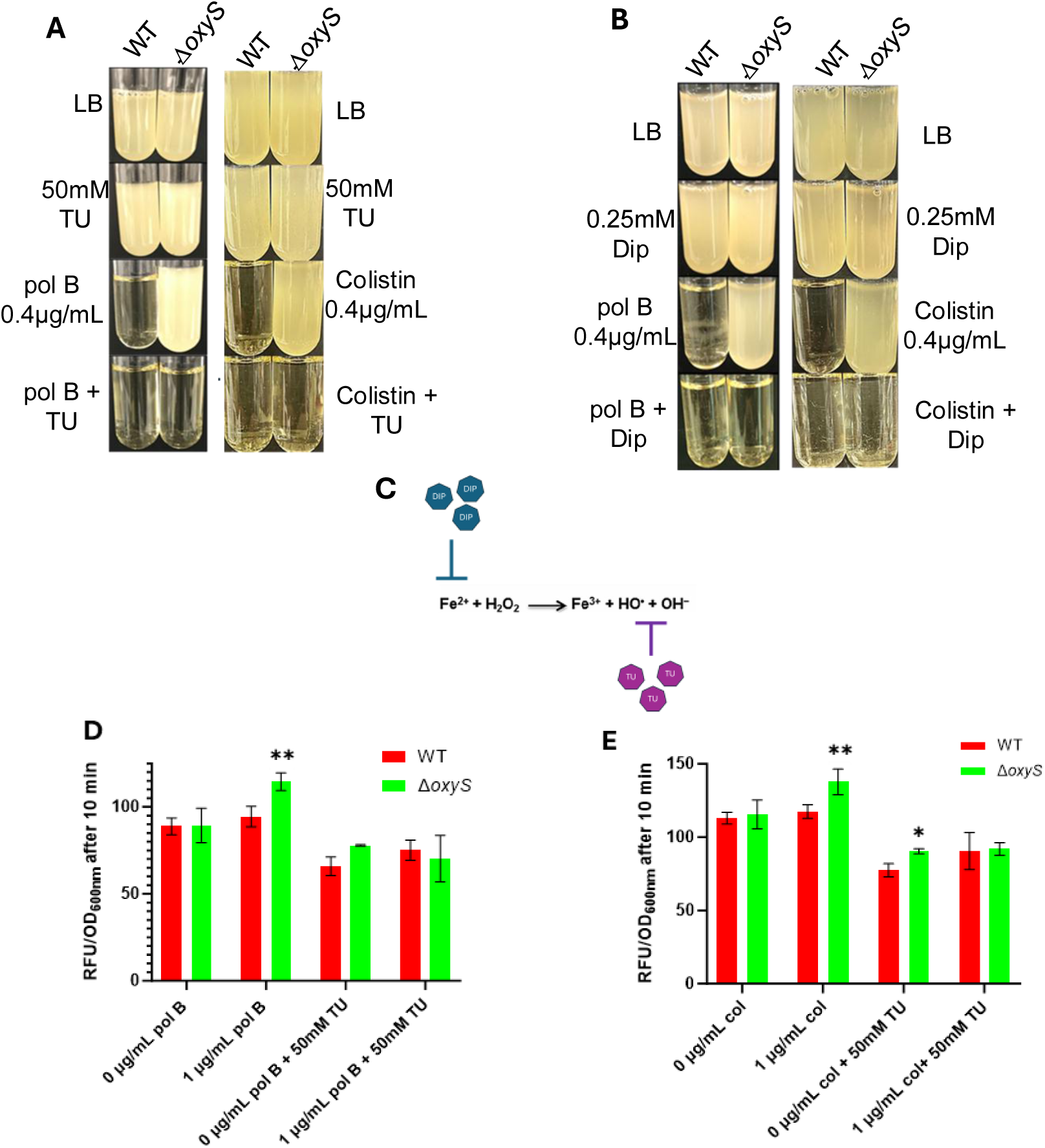
Resistance to polymyxin B or colistin in Δ*oxyS* mutant may depend on the accumulation of ROS. **(A)** Growth assay of WT and Δ*oxyS* mutant in the presence of polymyxin B (Pol B) or colistin (Col) at 0.4 µg/mL with or without thiourea (TU) at 50 mM. Cells were incubated at 37°C during 24 hours in LB medium (n=3). (**B)** Growth assay of WT and Δ*oxyS* mutant in the presence of polymyxin B or colistin at 0.4 µg/mL with or without 2.2 -dipyridyl (Dip) at 0.25 mM. Cells were incubated at 37°C during 24 hours in LB medium (n=3). (**C)** Mechanism of action of antioxidant thiourea (TU) and Dip. Thiourea (TU) prevents hydroxyl ions proliferation and Dip sequester Fe^2+^, thus preventing the Fenton reaction. **(D)** ROS quantification with HPF fluorescein dye (5 µM) on WT and Δ*oxyS* cells in the absence or in the presence of 1 µg/mL polymyxin B, with or without thiourea (TU) at 50 mM in M63 medium (n=3). **(E)** ROS quantification with HPF fluorescein dye (5 µM) on WT and Δ*oxyS* cells in the absence or in the presence of 1 µg/mL colistin, with or without thiourea (TU) at 50 mM in M63 medium (n=3). Statistical analysis were performed using Two-way ANOVA: Tukey s multiple comparisons test.

### Characterizing mutations from Δ*oxyS* variants resistant to polymyxin B and colistin

Results in Fig. 3 and 4 suggested that our Δ*oxyS* cells growing in the presence of polymyxin B or colistin had acquired mutation(s) conferring resistance to these antibiotics. To investigate this, we sequenced the genome of several Δ*oxyS* variants resistant to polymyxin B or colistin (Fig. 5A). We first performed whole-genome sequencing (WGS) on Δ*oxyS* variants resistant to polymyxin B (PolB^R^).

**Figure 5.**
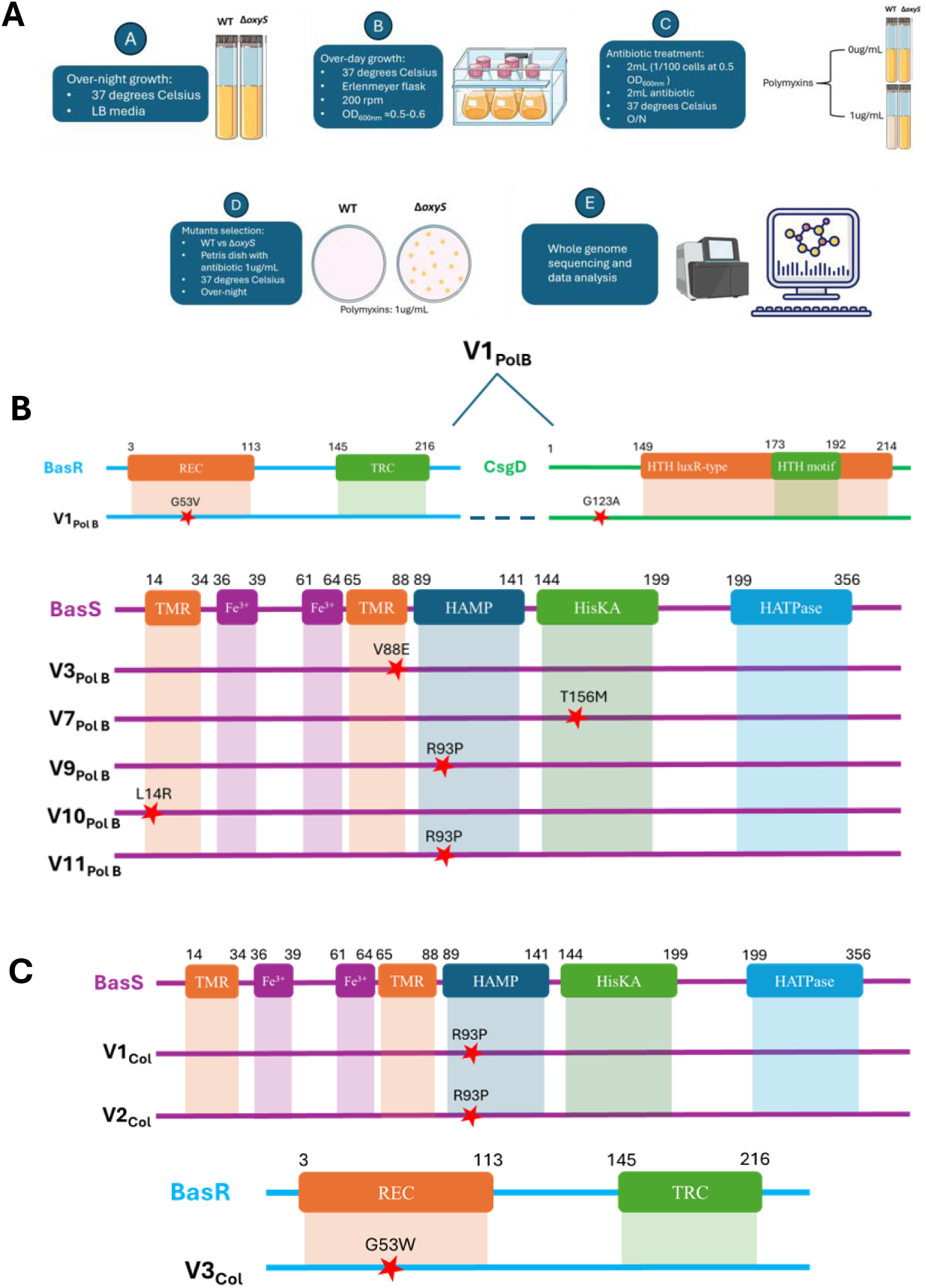
Characterizing mutations from Δ*oxyS* variants resistant to polymyxin B and colistin. **(A)** Schematic pathway for generating Δ*oxyS* variants resistant to polymyxin B (Δ*oxyS* PolB^R^) or colistin (Δ*oxyS* Col^R^). **(B)** Δ*oxyS* PolB^R^ *v*ariants and its mutations in amino acids sequence for BasR, BasS, and CsgD proteins. **(C)** Δ*oxyS* Col^R^ *v*ariants and its mutations in amino acids sequence for BasR and BasS Domains in BasR : REC : Receiver domain; TRC : Transcriptional regulatory protein, C terminal. Domains in BasS; TMR: Transmembrane regions; HAMP: Histidine kinases, Adenylyl cyclases, Methyl binding proteins, Phosphatases domain; HisKA : Kinase A domain; HATPases: Histidine kinase-like ATPases domain; Fe(III) : Conserved, Fe(III) binding motif. Domains in CsgD : HTH luxR type : Conserved region; HTH motif : DNA binding region.

Results in Fig. 5B show that one of these variants (V1) carried missense mutations, leading to a single amino acid substitution, in two different genes, *basR* and *csgD,* both encoding transcription factors. While BasR is a response regulator for BasS/R two-component system (TCS), promoting resistance to polymyxin and adaptation to neutrophils [35], the CsgD protein was previously shown to regulate biofilm formation [36]. WSG shows that the mutation in BasR (G53V) was in the regulatory domain region (REC), whereas the mutation in CsgD (G123A) was outside any known regulatory domains of the protein (Fig 5B, V1).

We further sequenced 5 more Δ*oxyS* variants resistant to polymyxin B and found missense mutations in a single gene, *basS,* which encodes an Fe-sensor of the BasS/BasR TCS. These 5 mutations demonstrated amino acids substitution in different domains of the BasS protein. For variants V3 (V88E) and V10 (L14R), mutations in amino acids were in different transmembrane domains (Fig. 5B). For variants V9 (R93P) and V11 (R93P), both variants share the same mutation in an amino acid located in the HAMP domain, involved in the regulation of BasS phosphorylation (Fig. 5B). The last variant resistant for polymyxin B, V7 (T156M), was mutated in the HisKA domain (Fig. 5B).

Moreover, we isolated and sequenced 3 Δ*oxyS* Col^R^ variants (Fig. 5C) and found two of them, V1 (R93P) and V2 (R93P) had substitution in the same amino acid as the V9 variant (HAMP domain of BasS). The last variant, V3 (G53W), was mutated in the regulatory region (REC) domain of the BasR protein (Fig. 5C), like V1. These results suggest that our Δ*oxyS* mutant can effectively generate mutations in genes previously shown to be involved in resistance to polymyxin B and colistin and potential new genes such as *csgD*.

### MepS protein expression promotes resistance to polymyxin B and colistin

Our results suggested that MepS could be involved in the development of polymyxin B and colistin resistance observed in Δ*oxyS* background (see Fig. 3B to 3E), suggesting a requirement for the peptidoglycan endopeptidase in the resistance mechanism. We wanted to examine the specific impact of MepS in the context of resistance against polymyxin B and colistin in our Δ*oxyS* variants. To address this, we introduced the Δ*mepS* allele in Δ*oxyS* variants resistant to polymyxin B (Δ*oxyS* PolB^R^) or colistin (Δ*oxyS* Col^R^). These PolB^R^ and Col^R^ variants were selected from experiments in Fig. 3B and Fig. 3C, respectively. Then, we performed a minimal inhibitory concentration (MIC) assay to assess the resistance of either Δ*oxyS* PolB^R^ Δ*mepS* variants or Δ*oxyS* Col^R^ Δ*mepS* variants. First, our results show that Δ*oxyS* PolB^R^ Δ*mepS* background were generally less resistant compared to Δ*oxyS* PolB^R^ variants when grown in the presence of polymyxin B (Fig 6A). Indeed, for the polymyxin B antibiotic the MIC decreased from 8 to 6 µg/mL for variant 1 (V1) and V3, from 6 to 2 µg/mL for V7, and from 8 to 4 µg/mL for V9. In contrast, V10 showed no change, with the MIC remaining at 6 µg/mL, despite the absence of the functional *mepS* gene (Fig 6A).

**Figure 6.**
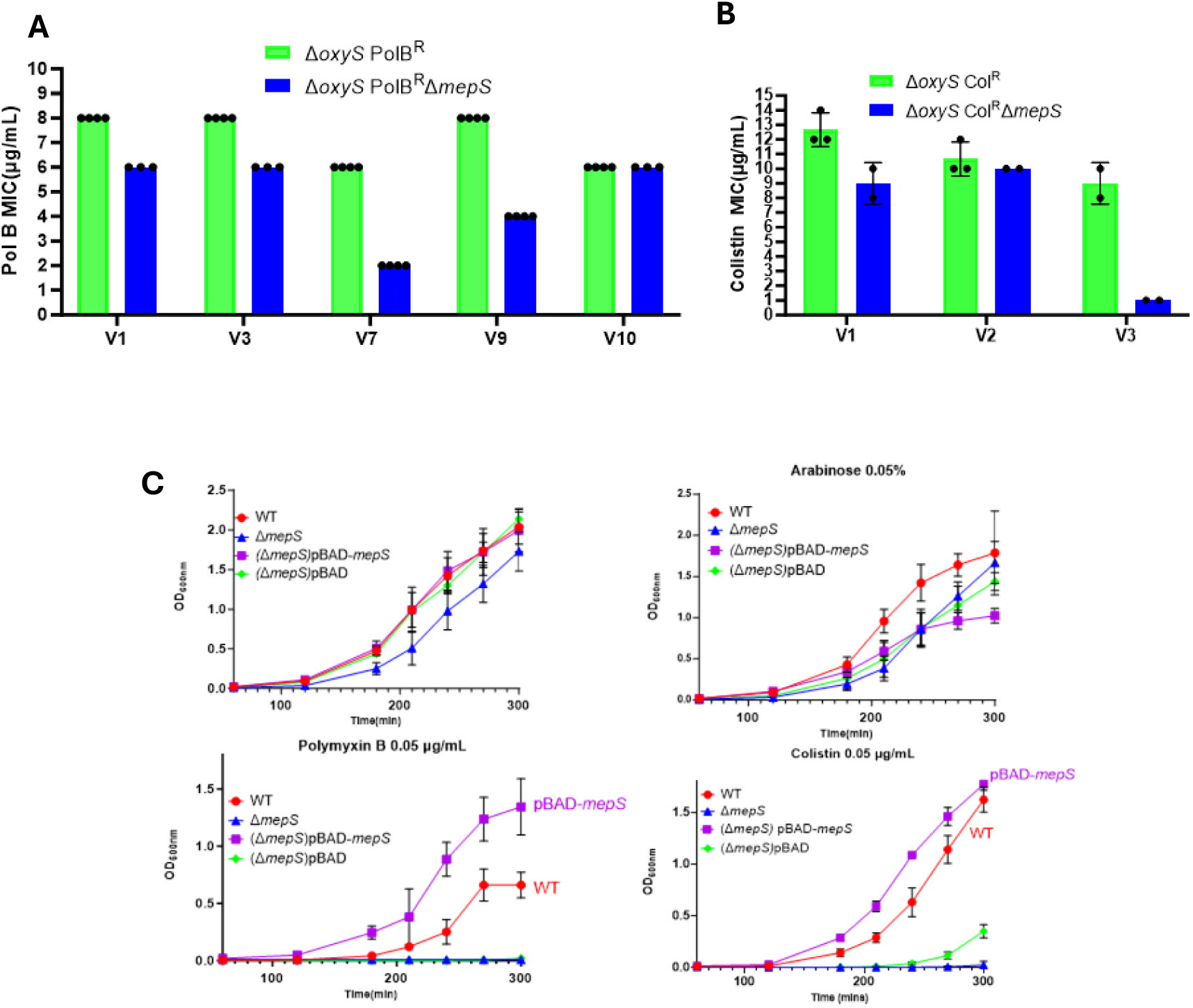
MepS protein contribution to polymyxin B or colistin resistance. **(A)** Minimal Inhibitory Concentration (MIC) of Δ*oxyS* PolB^R^ variant (green) and MIC of Δ*oxyS* PolB^R^ Δ*mepS* construct (blue). (**B**) MIC of Δ*oxyS* Col^R^ variant (green) and MIC of Δ*oxyS* Col^R^ Δ*mepS* construct (blue). (**C**) Growth curve of WT, Δ*mepS*, Δ*mepS+*pBAD-*mepS*Δ*mepS+*pBAD strains at 37°C LB (with or without polymyxin B or colistin antibiotics at 0.05µg/mL or arabinose at 0.05%) (n=3).

We also performed a MIC assay to assess the resistance to colistin of Δ*oxyS* Col^R^ Δ*mepS* background compared to the original Δ*oxyS* Col^R^ variants (Fig 6B). Similar to the results above, the MIC of colistin decreased slightly for V1 (from ∼12 to 10 µg/mL) and V2 (from ∼11 to 10 µg/mL), while a marked drop was observed for V3 (from 10 to 1 µg/mL) (Fig. 6B). These results suggest that Δ*oxyS* PolB^R^ or Δ*oxyS* Col^R^ variants generally rely on the presence of active MepS to exhibit full resistance to polymyxin B and colistin.

Next, we tested whether the overexpression of the MepS endopeptidase protein could also have an impact on the resistance to polymyxin B and colistin. To perform this experiment, we used a pBAD-*mepS* plasmid for the expression of the MepS protein and a pBAD empty vector as control (Fig. 6C). In these experiments, induction of plasmid pBAD-*mepS* with arabinose was omitted as we noticed a detrimental effect of the pBAD-*mepS* construct on growth compared to the empty pBAD vector (Fig 6C), suggesting that too much MepS protein could be toxic for the cells [59]. Next, we performed a growth curve with our strains (WT, Δ*mepS*, Δ*mepS* + pBAD-*mepS*, and Δ*mepS* + pBAD as control) in the presence of polymyxin B or colistin at 0.05 µg/mL. We noticed that the Δ*mepS* and the control Δ*mepS* + pBAD could not grow in presence of polymyxin B or colistin compared to the WT and Δ*mepS* + pBAD-*mepS* (Fig. 6C, 0.05 µg/mL). However, the Δ*mepS* + pBAD-*mepS* was able to grow better than the WT in presence of antibiotics at 0.05 µg/mL. These results suggest that the presence of MepS can help cells grow in the presence of polymyxin B or colistin.

### Absence of MepS protein may reduce the mutation rate in *E. coli*

Our results suggest that the repression of MepS by the sRNA OxyS could influence the mutation frequency and the emergence of resistance to antibiotics (Fig. 3 and Fig. 4). To investigate mutation frequencies across our various genetic backgrounds, we measured the mutation rate with a rifampicin assay, as previously described [18]. A rifampicin concentration of 100 µg/mL was used for the mutation frequency assay, as this concentration inhibits the growth of non-resistant cells (Fig supp 5A). To perform this assay, we grew our strains (WT, Δ*mepS*, Δ*oxyS*, and Δ*mepS-*Δ*oxyS*) in the presence of increasing concentrations of H_2_O_2_ (0 and 300 µM), to induce oxidative stress, and measured the development of resistance to rifampicin. First, during growth conditions with 0 µM H_2_O_2_, we did not observe any significant difference in the mutation rate between our four strains (Fig. 7A). In contrast, the rifampicin assays performed in the presence of 300 µM H₂O₂ revealed distinct mutation rates across strains (Fig. 7A). We observed that the mutant Δ*oxyS* had a higher mutation rate among all strains (Fig. 7A). This phenotype of the Δ*oxyS* mutant was also observed in a previous study [18]. We also found that the Δ*mepS* mutant displays a reduced mutation rate compared to the WT background. Moreover, the double mutant Δ*mepS*-Δ*oxyS* exhibits a mutation rate like the WT, indicating that the absence of *mepS* suppresses the mutator phenotype initially observed in the Δ*oxyS* background (Fig. 7A). These results may suggest that the absence of MepS may have a potential anti-mutator effect during oxidative stress in *E. coli*. We next assessed whether the mutation frequency in the presence of polymyxin B or colistin correlated with the phenotypes previously observed (Fig 3). The Δ*oxyS* mutant displayed an increased mutation frequency in the presence of both polymyxin B (Fig 7B) and colistin (Fig 7C) at 1 µg/mL compared with the WT strain. In contrast, the Δ*mepS* mutant showed a reduced mutation frequency under polymyxin B exposure (Fig 7B) and a frequency similar to the WT strain in the presence of colistin (Fig 7C). Notably, the absence of *mepS* in the Δ*oxyS*-Δ*mepS* double mutant decreased the mutation frequency for both antibiotics like the WT, suggesting that absence of MepS may contribute to limit emergence of resistance for polymyxins antibiotics.

**Figure 7.**
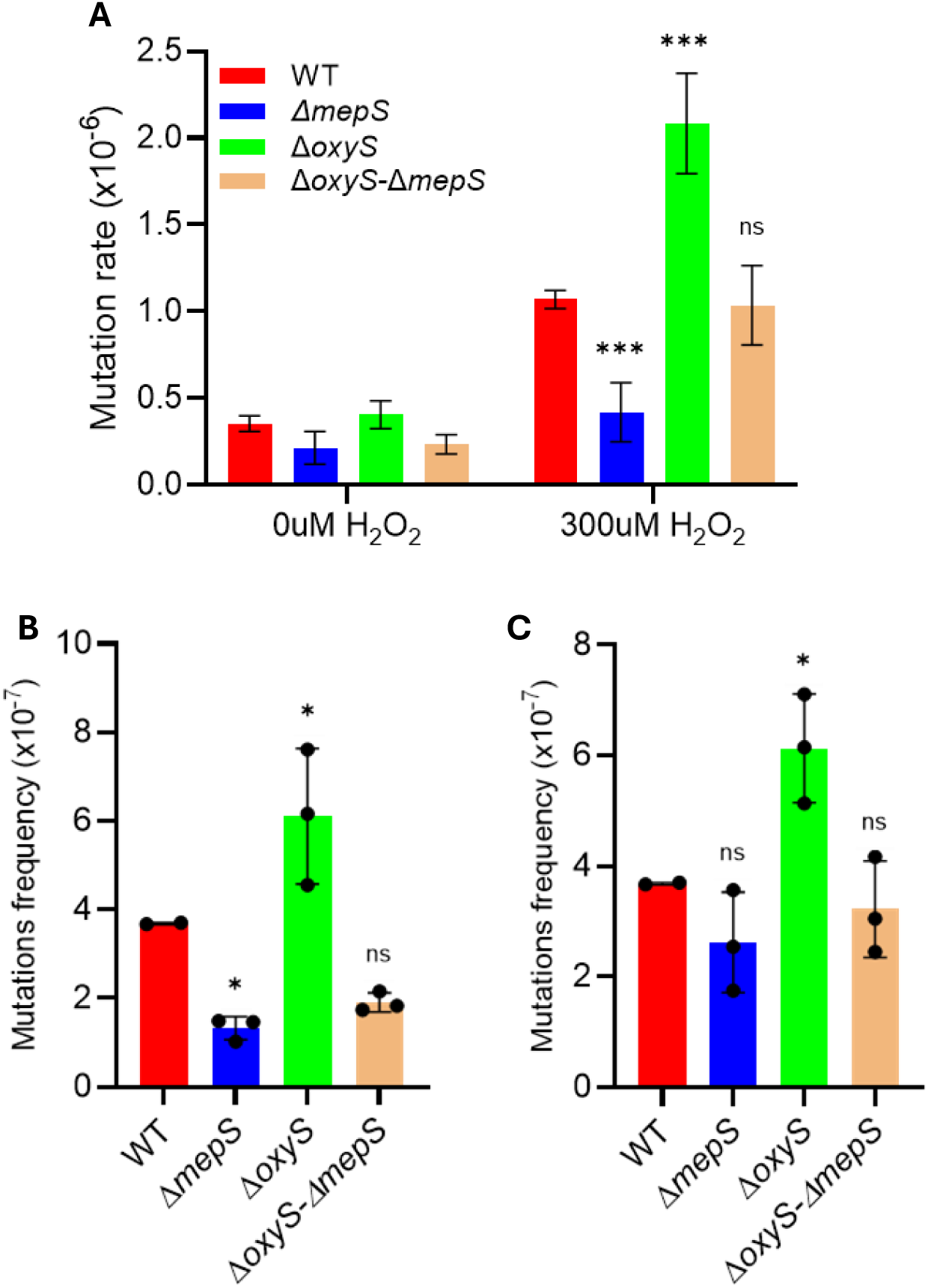
Absence of MepS could reduce the mutation rate in *E. coli*. Mutation rates of WT, Δ*mepS*, Δ*oxyS*, and Δ*mepS-*Δ*oxyS* cells grown in LB and plated on LB agar supplemented with 100 µg/mL rifampicin following exposure to **(A)** 0 µM H₂O₂ and 300 µM H₂O₂. **(B)** Mutation frequency of WT, Δ*mepS*, Δ*oxyS*, and Δ*mepS-*Δ*oxyS* cells grown in LB and plated on LB agar supplemented with 1 µg/mL of polymyxin B. **(C)** Mutation frequency of WT, Δ*mepS*, Δ*oxyS*, and Δ*mepS-*Δ*oxyS* cells grown in LB and plated on LB agar supplemented with 1 µg/mL of colistin. Mutation is expressed as the number of colonies on LB With antibiotics plates relative to the total CFU growing on LB. Data represents at least 3 independent experiments. Statistical analysis were performed using two-way ANOVA : Tukey s multiple comparaison test **(A)** and one-way ANOVA : Dunnett s multiple comparaison test **(B)** and **(C)**

**Figure 8.**
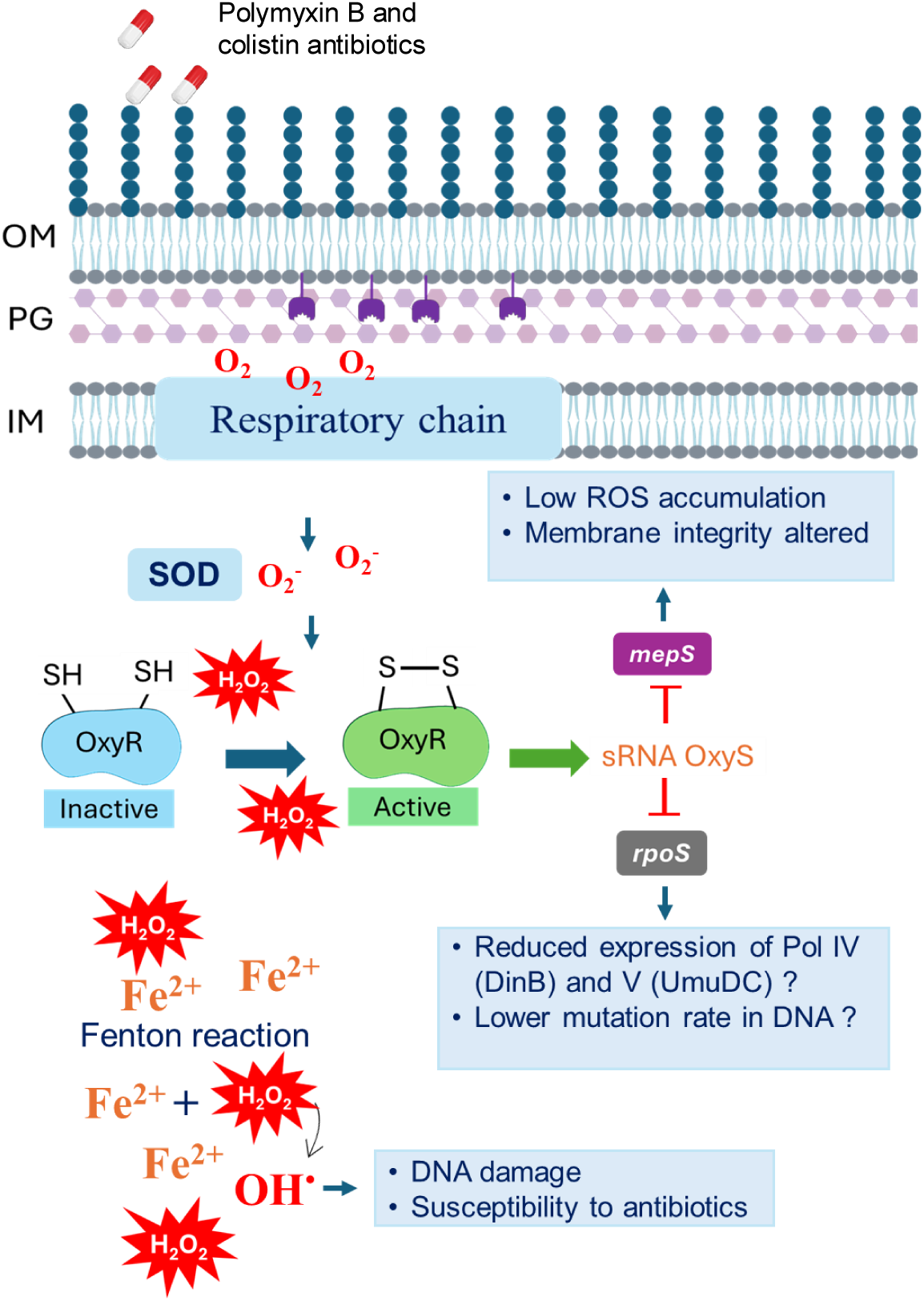
Schematic functions of the sRNA OxyS and MepS endopeptidase during oxidative stress induced by polymyxin B and colistin antibiotics. Polymyxin B and colistin treatment may disrupts *E. coli* respiratory metabolism, promoting ROS accumulation. Both ROS-derived H₂O₂ and Fe fuel the Fenton reaction, leading to DNA damage and activation of OxyR (via H₂O₂), which induces expression of the sRNA OxyS. OxyS represses *mepS* translation, increasing membrane fragility. Decreased MepS protein, may potentially influence the level of ROS in bacteria. Concurrently, OxyS represses *rpoS*, limiting the expression of mutagenic DNA polymerases Pol IV and Pol V. This dual action of OxyS, limiting both mutagenesis and cell wall robustness, creates a situation where *E. coli* becomes more susceptible to polymyxin B and colistin antibiotics.

## DISCUSSION

In this study, we investigated the potential phenotypes resulting from interaction between the sRNA OxyS and the target mRNA *mepS* in bacterial cells during oxidative stress. Our results indicated that Δ*oxyS* mutant promoted polymyxin B and E resistance compared to a wild-type strain, suggesting a function for OxyS in preventing resistance to these antibiotics. The data also suggested a potential role for *mepS* gene in the emergence of resistance to polymyxin B and E and in the maintenance of the resistance phenotype. While the Δ*oxyS* mutant strain could develop resistance to polymyxin B and colistin in our growth conditions, the double mutant Δ*oxyS-*Δ*mepS* remained sensitive to both antibiotics (Fig 3). Moreover, when we used OxySmut, a mutant version unable to bind its target mRNA *mepS*, the resistance phenotype in the presence of polymyxin B was similar to the Δ*oxyS* mutant (see supp Fig 4). These results could suggest that the absence of *mepS* may limit the emergence of resistance in the Δ*oxyS*-Δ*mepS* background and the repression of sRNA OxyS is important to limit emergence of resistance as we observed in *oxyS*mut strain version. Moreover, when the MepS protein was overproduced by a plasmid, the strains displayed better growth than the WT in the presence of polymyxin B and colistin antibiotics (Fig 6). These results may suggest that the peptidoglycan synthesis activity of MepS could potentially be involved in the phenotype of resistance. Indeed, the absence of MepS could increase the vulnerability of the bacterial cell wall to polymyxin B and colistin (Fig. 2A to C). When we deleted the *mepS* gene from Δ*oxyS* PolB^R^ variants or Δ*oxyS* Col^R^ variants, the resulting Δ*oxyS* PolB^R^ Δ*mepS* and Δ*oxyS* Col^R^ Δ*mepS* constructs often exhibited reduced resistance to both antibiotics (Fig 6A-B).

Although the absence of MepS could make cells more vulnerable to envelope stress (Fig 2), the Δ*mepS* mutant displayed sensitivity to polymyxin B and colistin similar to that of the WT under our conditions (Fig 3B, C, D, and E). We therefore favor the hypothesis that the absence of *mepS* could limit the emergence of resistant mutants, potentially by reducing the emergence of mutation rate in DNA or affecting cellular processes required for resistance acquisition.

Indeed, it is also possible that a repression of MepS could limit mutation rates in DNA (Fig. 7A), which may rely on ROS accumulation for the development of polymyxin resistance (Fig. 4). A previous study showed that sublethal antibiotic exposure can promote resistance development by inducing oxidative stress and DNA damage, thereby increasing mutation rates and facilitating the emergence of resistance trough several days [14]. Consistent with this, resistance in the Δ*oxyS* mutant emerged after approximately 10 h of growth for both polymyxin B and colistin antibiotics, potentially reflecting the emergence of a resistant subpopulation (Fig 3D-E) and not in the double mutant Δ*oxyS*-*mepS*. Also, the mutation frequency assay further supported these findings, as the Δ*oxyS* mutant displayed a higher mutation frequency than the WT, suggesting an increased potential for the emergence of resistant subpopulations (Fig 7). The WGS confirms mutations in Δ*oxyS* PolB^R^ and Δ*oxyS* Col^R^ variants in the BasSR (PmrAB) two-component system frequently associated with polymyxin resistance in clinical isolates of *E. coli*, *Salmonella enterica*, *K. pneumoniae*, and *Acinetobacter* spp. [45] [46]. Our data also indicated mutation in *csgD*, encoding the CsgD transcription factor, which is involved in biofilm formation [36] [48]. To our knowledge, this is the first report on a mutation in *csgD* associated to polymyxin resistance. Our results suggest that resistance emergence in the Δ*oxyS* mutant may depends on ROS accumulation and intracellular iron (see Fig. 4). Indeed, treatment with thiourea or the iron chelator 2,2′-dipyridyl prevented the emergence of polymyxin B and colistin resistance, likely by reducing ROS mediated DNA mutation.

The sRNA OxyS was previously shown to repress *rpoS* mRNA during oxidative stress [41], which could also partly explain our phenotype in mutant Δ*oxyS*. RpoS is a general stress response transcription factor and regulates the expression of DNA repair genes and induces expression of error-prone DNA polymerases such as Pol IV (DinB) and Pol II [42] [43]. These DNA polymerases may lead to an increase of missense mutations within the nucleotide sequence, facilitating adaptation through mutations in key genes [44]. The absence of OxyS could potentially enhance *rpoS* expression during polymyxin B and colistin treatments known to induce oxidative stress [34]. This could represent a potential adaptation mechanism to the selective pressure exerted by polymyxin B and colistin in our Δ*oxyS* mutants. Also, absence of OxyS could have an impact on Fe-S clusters. Indeed, OxyS can promote the expression of *iscRSUA* operon, which converts oxidative-stress inducing Fe^2+^ into more stable Fe-S clusters, potentially alleviating the ROS-producing Fenton reaction [23].

According to Barshishat et al [20], the sRNA OxyS can indirectly help to repair DNA damage by repressing the *nusG* mRNA, which stops the bacterial cell cycle [31]. Concurrently with *nusG* repression, the sRNA OxyS could slow down cell division during oxidative stress by decreasing the level of MepS proteins in bacterial cell (Fig 1D-E) by affecting peptidoglycan synthesis [62]. This may also help bacteria repair intracellular damage caused by ROS and effectively decrease mutations in DNA.

To explore whether reduced MepS levels may influence DNA mutagenesis during stress we characterized the effect of a Δ*mepS* mutant under various conditions. Our results with rifampicin assay showed that the Δ*mepS* mutant exhibited a lower mutation rate than the wild-type strain under oxidative stress induced by 300 µM of H_2_O_2_ (Fig.7A). At 0 µM H₂O₂, no significant difference was observed between the strains, suggesting that the mutation frequency is not simply due to an intrinsic growth or sensitivity but rather is associated with the oxidative stress conditions that promote the emergence of rifampicin-resistant mutations. The absence of *mepS* also reduced the mutation rate in the double mutant Δ*oxyS*Δ*mepS* compared with the Δ*oxyS* single mutant (Fig. 7A). Consistent with these findings, polymyxin B and colistin selection assays showed that Δ*mepS* mutants developed reduced resistance compared with the wild-type strain (Fig. 7B and C, respectively). Also, deletion of *mepS* attenuated the enhanced resistance phenotype against polymyxin B and colistin associated with the Δ*oxyS* mutant (Fig 7B and C, respectively). Together, these results seem to indicate that absence of MepS could contribute to diminishing mutation in DNA. potentially by reducing oxidative stress.

Decreased MepS levels were shown to alter peptidoglycan synthesis [28]. Thus, it could be possible that decreased MepS could potentially affect the respiratory chain anchored to the inner membrane and indirectly lead to a reduced production of ROS [52] [53] [54]. We also observed that the Δ*mepS* mutant displays reduced cell count compared with the wild-type strain (Fig. S6). The decreased cell counts in the Δ*mepS* mutant could suggest a repressed DNA replication compared to WT, which may limit the proliferation of cells carrying DNA damage.

In *E. coli*, MepS activity is functionally redundant with peptidoglycan endopeptidases MepH and MepM to facilitate peptidoglycan synthesis during cell elongation [49] [29]. Despite the presence of MepH and MepM proteins, the double mutant strain Δ*oxyS-*Δ*mepS* could not develop resistance to polymyxins like the Δ*oxyS* mutant. This observation may suggest a specific role for the MepS protein in the presence of polymyxins. Hyoung Park et al. [50] showed that these three proteins have different role depending on growth conditions. Indeed, Δ*mepS* mutants are specifically sensitive to EDTA, likely because MepM inhibition cannot be compensated for in the absence of MepS [50].

Taken together, our results support that during oxidative stress, the sRNA OxyS represses *mepS* translation, reducing MepS protein levels in bacteria cell (Fig 1D-E). By characterizing the Δ*mepS* mutant under oxidative stress conditions, we found that the absence of MepS decreases mutation frequencies, suggesting that a potential OxyS mediated repression of *mepS* mRNA during oxidative stress may contribute to limiting DNA mutagenesis. Although the precise molecular mechanism remains to be elucidated, our model provides a conceptual framework integrating peptidoglycan remodeling, oxidative stress response, and polymyxin resistance in *E. coli*. In perspective, we will explore the possibility that Δ*mepS* mutant has less intracellular ROS, which would prevent emergence of DNA mutations. Our future experiments will help determine whether peptidoglycan endopeptidase activity and ROS are related.

## MATERIALS AND METHODS

### Strain and Growth

*E. coli* K-12 MG1655 derivatives were used throughout this study. Strains generated by P1 transduction carried specific antibiotic markers. Unless stated otherwise, 50 µg/mL ampicillin was used for pNM12 and pBAD-OxyS plasmid selection. See Supplemental Table S1 for strains, plasmids, and Table S2 for oligonucleotide details. All strains were grown in LB medium unless stated otherwise. Overnight cultures were incubated at 37°C, with ampicillin (50 µg/mL) added for plasmid maintenance. Experiments were performed in 50 mL of LB medium in 250 mL flasks at 37°C, shaking at 220 rpm, with or without ampicillin. OxyS overexpression was induced with 0.1% arabinose at OD_600nm_ = 0.1. Endogenous OxyS was induced with H_2_O_2_ at various concentrations, starting at OD_600nm_ of 0.5. RNA and protein samples were collected at multiple time-points post-induction.

### Total RNA extraction

Cells were grown in 3 mL of LB medium with ampicillin, if necessary, overnight at 37°C. The following day, 50 μL of overnight cultures were inoculated at 37°C for experiments. Arabinose or H_2_O_2_ was added as indicated for the induction of OxyS after reaching an OD_600nm_ of 0.1 or 0.5, respectively. 600 μL of culture samples were collected at different time-points after OxyS induction. Total RNA was extracted using the hot phenol protocol described previously [55]. RNA samples were quantified using the Nanodrop Denovix (DS-11+ Spectrophotometer).

### OxyS DNA probe

The OxyS DNA probe was labeled with P32ɣ-ATP using T4 polynucleotide kinase (T4-PNK) from New England Biolabs. In a microtube, we prepared a mixture of 13 μL Millipore water, 2 μL of 10X PNK Buffer, 2 μL of 5 μM oligonucleotide, 2 μL of P32ɣ-ATP (6000Ci), and 1 μL of T4-PNK. The mixture was incubated at 37°C for 30 minutes, and the volume was then adjusted to 100 μL with Millipore water. The mixture was passed through Sephadex G-50 columns (5 minutes at 2500rpm) to purify the probe. 50 μL of the purified probe was used for overnight hybridization at 37°C. The remaining probe was stored at -20°C for future use.

### Northern blot

10 μg of total RNA were loaded on a 5% polyacrylamide gel (TBE, Urea) after heating samples at 90°C for 1 min. Electrophoresis was run in 1X TBE at 100 V for 1 h. RNAs were then electro-transferred onto a Hybond-XL membrane (Amersham Biosciences) in 0.5X TBE at 200 mA for 1 h. RNAs were UV cross-linked at 254 nm for 1 min. Pre-hybridization (30 min at 42°C) and hybridization were performed in 10 mL Church buffer (Church and Gilbert, 1983) with OxyS DNA probe, overnight at 42 °C. Membranes were washed twice (5 min in 2X SSC-0.1% SDS) and twice (15 min in 0.1X SSC-0.1% SDS) at 42°C. Finally, membranes were stored in phosphor cassettes overnight at room temperature and analyzed with a Typhoon Trio (GE Healthcare).

### Total protein extraction

Cells were cultured overnight at 37°C in 3 mL of LB with ampicillin (50 mg/mL, 1:1000 dilution, if needed). The next day, 50 μL of cultures were inoculated at 37°C for experiments (see strains and growth conditions above). Transcription of OxyS sRNA was induced with 0.1% arabinose or H_2_O_2_ at OD_600nm_ = 0.5. For protein extraction, 1 mL of culture was collected in Eppendorf tubes. The volume was adjusted based on OD_600nm_ using (OD_600nm_ / 0.5) to normalize protein quantity. Samples were centrifuged (1 min, 13,000 rpm), pellets were resuspended in 1 mL LB, then mixed with 110 μL cold 50% TCA and kept on ice. After vortexing for 5 seconds and 10 min incubation on ice, samples were centrifuged (10 min, 13,000 rpm, 4°C), washed twice with 500 μL ice-cold 80% acetone, and centrifuged again (5 min, 13,000 rpm, 4°C). Supernatants were discarded, and pellets were air-dried (1–2 min), then resuspended in 100 μL of 1X protein buffer (100 mM DTT) and stored at –20°C.

### Western blot

We used 5 μL of extracted total proteins for the western blot transfer. Samples were loaded onto a 12% SDS-PAGE gel in 1X running buffer and electrophoresed at 150V for 1 hour and 20 minutes. Protein samples were heated at 95°C for 3 minutes before loading. Protein transfer from the gel to a nitrocellulose membrane was performed by electro-transfer in 1X transfer buffer at 250 mA for 1 hour. After transfer, the western blot procedures were carried out according to [56]. The western blot results were analyzed using an Odyssey infrared imaging system on the Li-Cor Biosciences instrument, and quantification was performed using the Odyssey application software.

### β-galactosidase assay

Cells were cultured overnight at 37°C in 3 mL of LB medium with or without ampicillin (50 mg/mL, 1:1000 dilution). The next day, 50 μL of these cultures were incubated in 50 mL of LB with 50 ug/mL ampicillin at 37°C, shaking at 220 rpm. Expression of OxyS was induced by adding 0.1% arabinose at OD_600nm_ = 0.1. For endogenous OxyS expression, 300 μM H_2_O_2_ was added every 15 min after about 1 h of growth (OD_600nm_ = 0.1) until the end of the experiment. β-galactosidase activity was measured as in [57] using a SpectraMax 250 microplate reader. Specific activity was calculated as V.max/OD_600nm_.

### Lead Acetate probing assay

The lead acetate probing assay was conducted following previously established protocols [56]. In a concise overview, 0.1 μM of γ-32P radiolabeled *mepS* was incubated with or without 1 μM of sRNA OxyS for 15 minutes at 37°C, followed by treatment with 5 mM PbAc for 2 minutes. Controls were treated with H_2_O. The reactions were halted by the addition of 10 μl of Loading Buffer II (LBII: 95% formamide, 18 mM EDTA, 0.025% SDS, 0.025% xylene cyanol-bromophenol blue). Subsequently, samples were electrophoresed on polyacrylamide gels (8% acrylamide: bisacrylamide 19:1, 8 M urea) in TBE 1X at 38 W. The gels were dried, exposed to phosphor screens, and visualized using the Typhoon Trio (GE Healthcare) instrument.

### Growth curve assay

Overnight cultures were grown in LB medium at 37◦C. The next day, we calculated (0.5/ OD_600nm_ overnight) to obtain an OD_600nm_ of 0.5 in 1mL total. Then we put 100µl of OD_600nm nm_=0.5 in microplate with (100µl) or without antibiotics in LB medium. The cultures were incubated at 37◦C and OD_600nm_ was measured using BioTek EPOCH2T microplate reader.

### MIC assay

Overnight cultures were grown in LB medium at 37°C. The next day, the OD_600nm_ of the culture was measured and diluted to obtain 10^8^ cells/mL. Following this, we performed dilutions from 10^8^ cells/mL to 10^6^ cells/mL in LB medium and put 100µl of 10^6^ cells/mL in microplate with antibiotics (100 µl). The cultures were incubated at 37°C for 24 hours using BioTek EPOCH2T microplate reader. The MIC was defined as the lowest concentration of polymyxins antibiotics at which no visible bacterial growth was observed after growth, as determined by the absence of turbidity compared to the growth control, indicating a complete inhibition of bacterial proliferation under the tested conditions

### Growth assays in the presence of antibiotics

Cells were grown in 250 ml flasks at 220rpm shaking for approximately 3 hours at 37°C until the culture reached an OD_600nm_ of 0.5. We then diluted the cultures 1/100 in LB medium in sterile tubes containing antibiotics at 1 µg/mL, 0.5 µg/mL, 0.25 µg/mL, 0.125 µg/mL, 0.0625 µg/mL final concentration. The tubes were incubated at 37°C overnight.

### CPRG assay

We used strains derived from MG1655 with a functional endogenous *lacZ* gene. The CPRG assay was conducted with some modifications from previously established protocols [58]. All experiments were performed in LB0N medium (LB medium without NaCl). Isolated colonies were streaked on Petri dish containing 200 ug/mL of CPRG and incubated for 18 hours at 37°C. For the liquid assay in 96 wells microplates, we used CPRG at 30 ug/mL final concentration. Overnight cultures were inoculated at 37°C in LB medium. Cultures were washed in LB0N before using them for the assay. 100 μL of our culture (at OD_600nm_ =0.5) was incubated with 100 μL of CPRG 60 ug/mL to have a final CPRG concentration of 30 ug/mL. When CPRG and polymyxin B were used together for the liquid assay, we used 10 ug/mL final of CPRG.

### MS2 Affinity Purification

Affinity purifications were adapted from a previous report [56]. Cells were grown to an OD_600nm_ = 0.5, induced with 0.1% arabinose for 10 min, and then chilled on ice. Total RNA was extracted from 600 μL of culture (input), and the rest was processed for lysis in buffer A (20 mM Tris-HCl pH 8.0, 150 mM KCl, 1 mM MgCl₂, 1 mM DTT) using a French Press (430 psi, 3×). Lysates were centrifuged (17,000 g, 30 min, 4°C), and 20 μL was saved for protein input. Affinity steps were performed at 4°C using 75 μL amylose resin loaded with 100 pmol of MS2-MBP. Lysates were loaded, washed, and eluted with buffer A + 15 mM maltose. RNA was purified by phenol-chloroform and ethanol precipitation and proteins by acetone precipitation. RNA and protein were analyzed by northern and western blots, respectively. 50 mL of exponential and stationary phase cells (OD_600nm_ = 0.5 or 1.0,) were lysed (8000 psi, 4×) and processed on a shared column with 100 μL resin and 200 pmol MS2-MBP.

### DNA genomic extraction

We inoculated 5 mL of our Δ*oxyS* variants resistant to polymyxin B (Δ*oxyS* PolB^R^) or colistin (Δ*oxyS* Col^R^) and WT backgrounds in LB medium at 37°C overnight. The next day, we centrifuged the samples at 3500 rpm for 10 minutes and removed the supernatant. Pellets were resuspended in 1900 µL of TE buffer (Tris 10mM, EDTA 1mM), then divided into 4 Eppendorf tubes of 475 µL each. We added 30 µL of 10% SDS and 3 µL of 20 mg/mL proteinase K, then incubated for 1 hour at 37°C. Next, we added 500 µL of phenol (pH 8.0), chloroform-isoamyl, mixed by inversion for 5 minutes, and centrifuged at 13,000 rpm for 10 minutes. We recovered the upper phase and repeated the phenol-chloroform extraction three times, ending with a chloroform-only extraction. To precipitate the DNA, we added 1/10 volume of 3M NaAcetate at pH 5.4, and 2 volumes of 100% ethanol, then incubated at -20°C. After centrifugation at 4°C for 10 minutes, we removed the supernatant and washed the DNA with 500 µL of 70% ethanol and centrifuging for 2 minutes. After drying the pellet, the DNA was resuspended in 200 to 500 µL of 10 mM Tris buffer. Finally, we treated the sample with RNase (20 µg/mL) at 37°C for 30 minutes and quantified the extracted DNA.

### Mutant clone selection

Cells were incubated in 3 mL of LB medium at 37°C overnight. The following day, we transferred 50 µL of our culture in 50 mL of fresh LB medium in 250 mL flask at 37°C and shaking at 220 rpm until an OD_600nm_ =0.5 (about 3 hours). Then, cultures were diluted 1/100 in tubes with polymyxin B or colistin antibiotics at 1 ug/mL and incubated at 37°C over-night. The next day, the cultures were spread on a Petri dish containing antibiotics (1 µg/mL). The following day isolated colonies were streaked on new Petri dishes with polymyxin B and colistin antibiotics at 1 ug/mL and then conserved at -80°C until further use.

### Rifampicin mutagenesis assay

Overnight cultures were diluted 1/1000 into 50 mL of LB medium in 250 mL flasks and grown at 37 °C with shaking. At an OD₆₀₀ of 0.2, H_2_O_2_ was added at different concentrations (0, 100, 200, and 300 µM), and cultures were incubated overnight. The next day, 10 mL of each culture was harvested by centrifugation at 2,500 × g for 15 min. Cell pellets were washed with 1 mL of PBS 1X and resuspended, then diluted 1/10 in PBS 1X for plating on LB agar supplemented with 100 µg/mL rifampicin. After overnight incubation at 37 °C, colonies were counted and the mutation rate was calculated as the ratio of rifampicin-resistant CFUs to total CFUs. Total CFUs were determined by serial dilution in PBS 1X and plating on LB agar.

### ROS quantification

Overnight cultures were grown in LB medium at 37◦C. The next day, cultures were washed in M63 medium and diluted to an OD_600nm_ = 0.5 in 1mL total. Then, cultures were transferred into a black 96 well microplate containing 5 µM of HPF, with or without antibiotics, with or without thiourea at 50 mM. The cultures were incubated at 37◦C and ROS level was measured using Synergy HTX microplate reader (Emission 485/20 and Excitation 528/20).

### Whole Genomic Sequencing

Genomic DNA samples were purified using AMPure beads at a 1× ratio. A total of 200 ng of purified DNA was used to prepare the sequencing library with the Rapid Barcoding Kit (SQK-RBK114.94, Oxford Nanopore Technologies), following the manufacturer’s instructions. The final library was loaded onto a PromethION flow cell for sequencing. Basecalling was performed using the Super Accuracy model in Dorado. The resulting raw sequencing data (FASTQ files) were analyzed using the *wf-bacterial-genomes* pipeline from Epi2me Labs (Nanopore) (https://github.com/epi2me-labs/wf-bacterial-genomes), with default parameters.

### Whole Genomic Sequencing Analysis

Adapter and quality trimming of fastq files were performed with Cutadapt v.2.8. Mapping to reference genome *E. coli* strain k12 substr mg 1665 gca 000005845 version ASM584v2.dna.toplevel was done with bwa mem v.0.7.17-r1188 with arguments K set 100000000 and using soft clipping for supplementary alignments (-Y). All the samples had on average more than 99% reads aligned. Duplicate reads were identified with gatk v.4.2.6.1 MarkDuplicates with arguments validation stringency set to silent, optical duplicate pixel distance set to 2500, clear dt set to false, assume sort order set to false and add pg tag to reads set to false. Aligned BAM files were sorted with gatk v.4.2.6.1 Sort Sam with arguments sort order to coordinate, create index to true and max records in ram to 30000. Read groups were added with gatk v.4.2.6.1 AddOrReplaceReadGroups with arguments RGLB to LaneX, RGPU to NONE, RGSM to sample name and RGPL to nanopore. Variant calling was done with gatk v.4.2.6.1 HaplotypeCaller with arguments sample ploidy to 1. Quality variant filtration was done with gatk v.4.2.6.1 VariantFiltration with arguments filter name to FS and lowQualFilter, filter to FS > 30, cluster window size to 10, cluster size to 3, missing values evaluate as failing ON, filter expression to QUAL < 30, QD < 2.0, FS > 60.0 and DP < 10. Variant annotation was performed with Variant Effect Predictor (VEP) at EnsemblBateria (https://bacteria.ensembl.org/Escherichia_coli_str_k_12_substr_mg1655_gca_000005845/Tools/VEP) using the same reference genome we used for mapping.

## ACKNOWLEDGMENTS

This work was funded by an operating grant BMB389354 from the Canadian Institutes of Health Research (CIHR) to EM.

