## Supplemental figures S1-S7 and Table 1-2 for "Peptidoglycan remodeling prevents antibiotic resistance during oxidative stress"

This file includes :

Supplemental figures S1 to S6 and Table S1 to S2

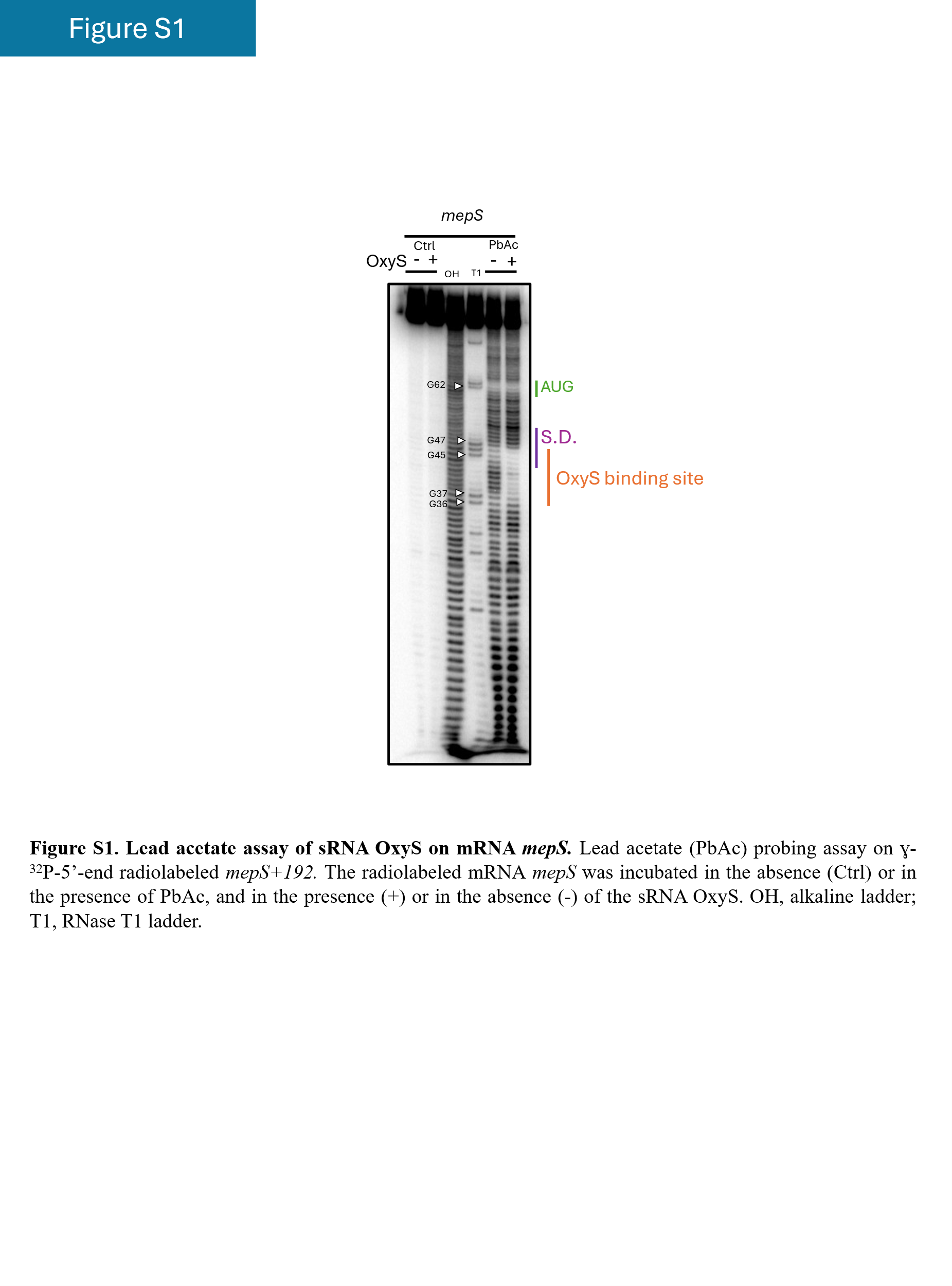

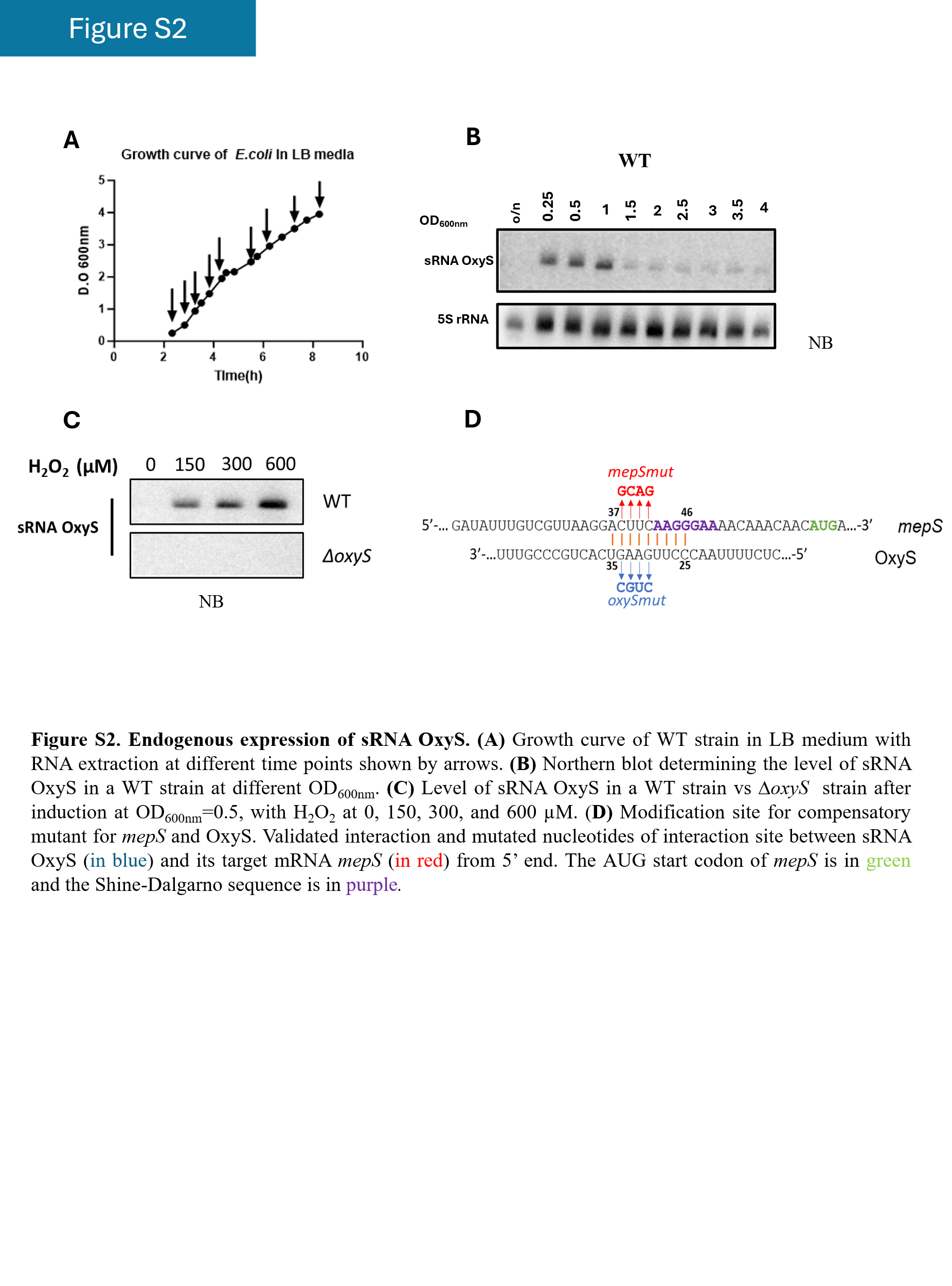

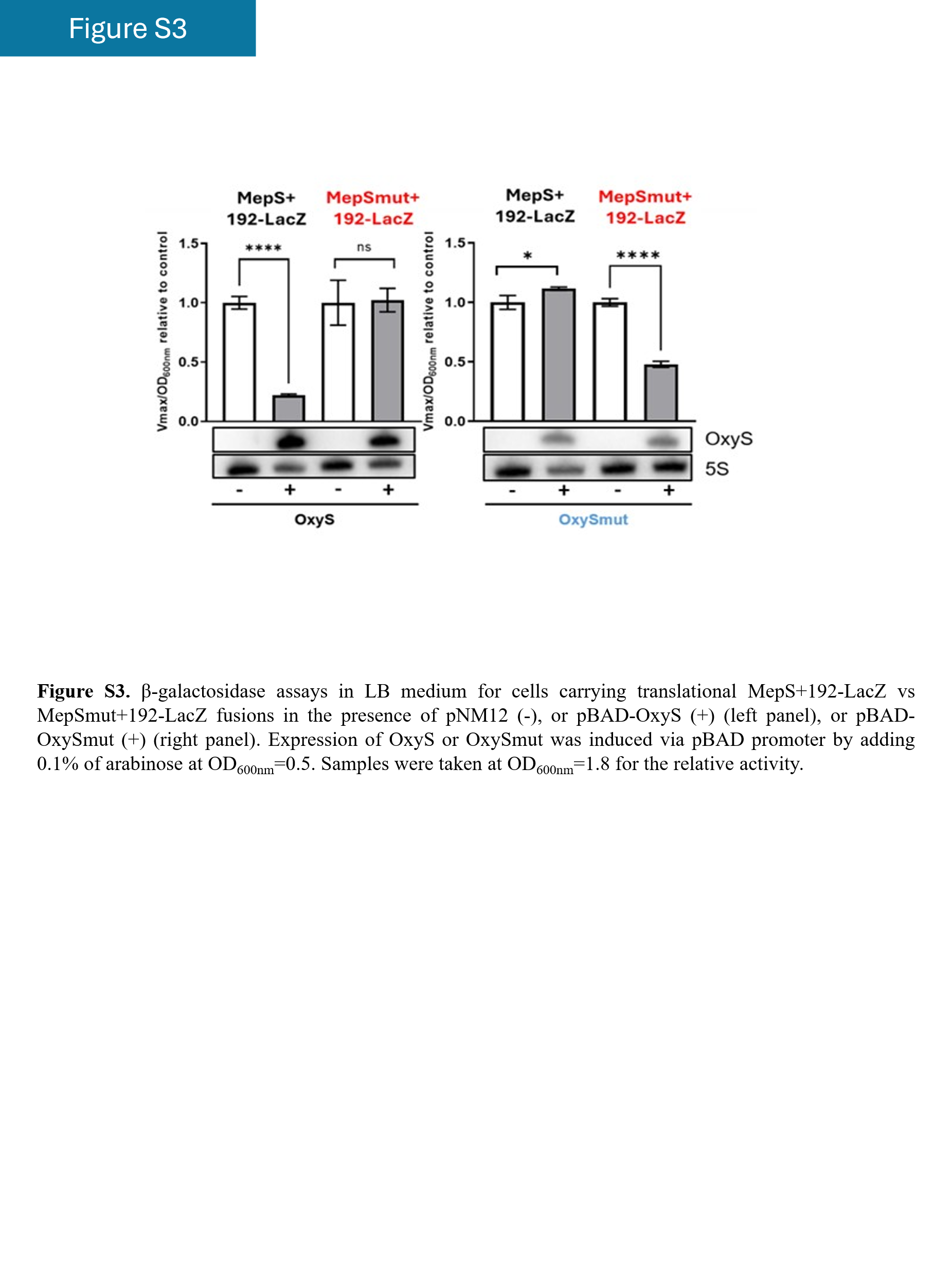

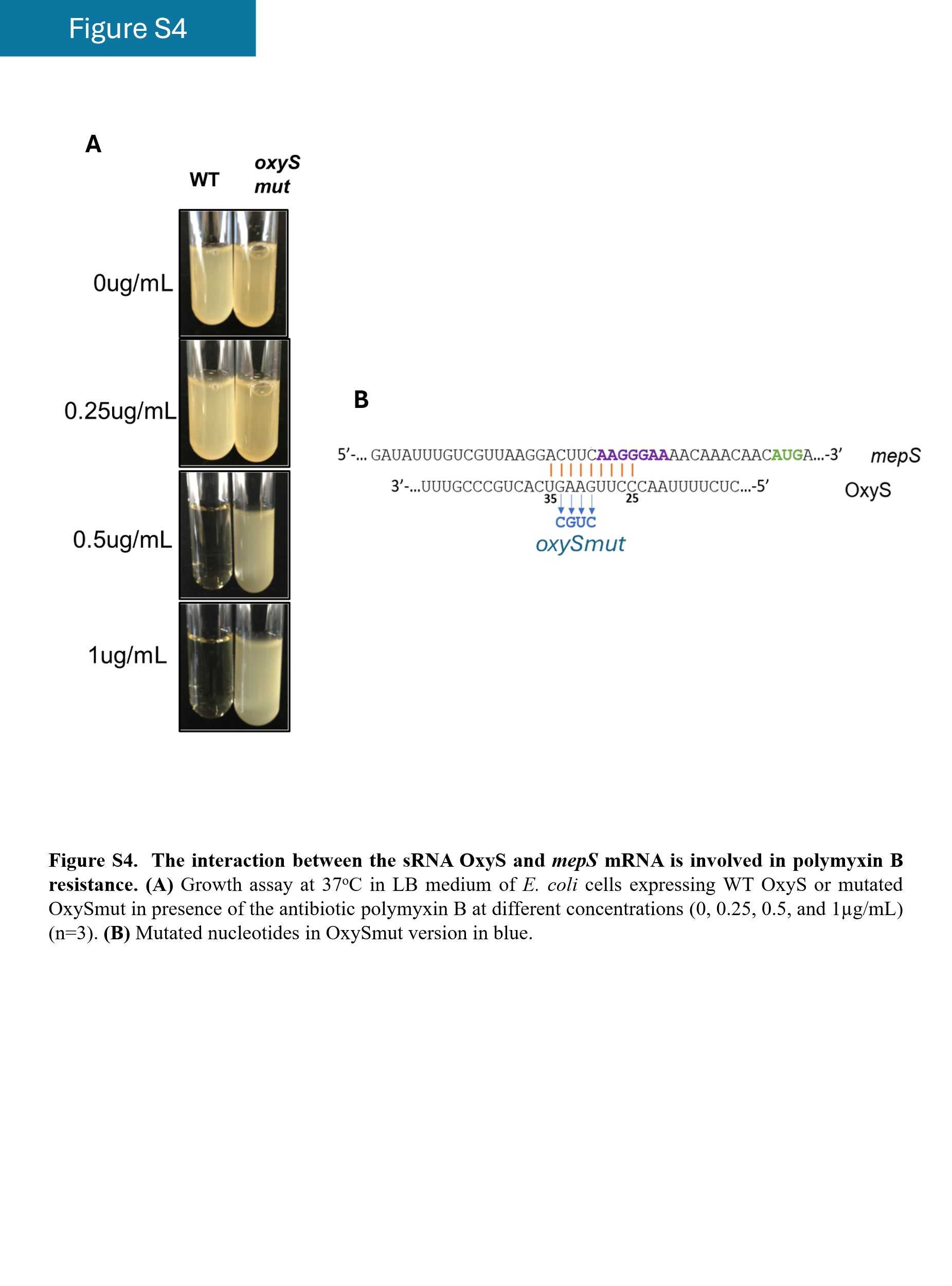

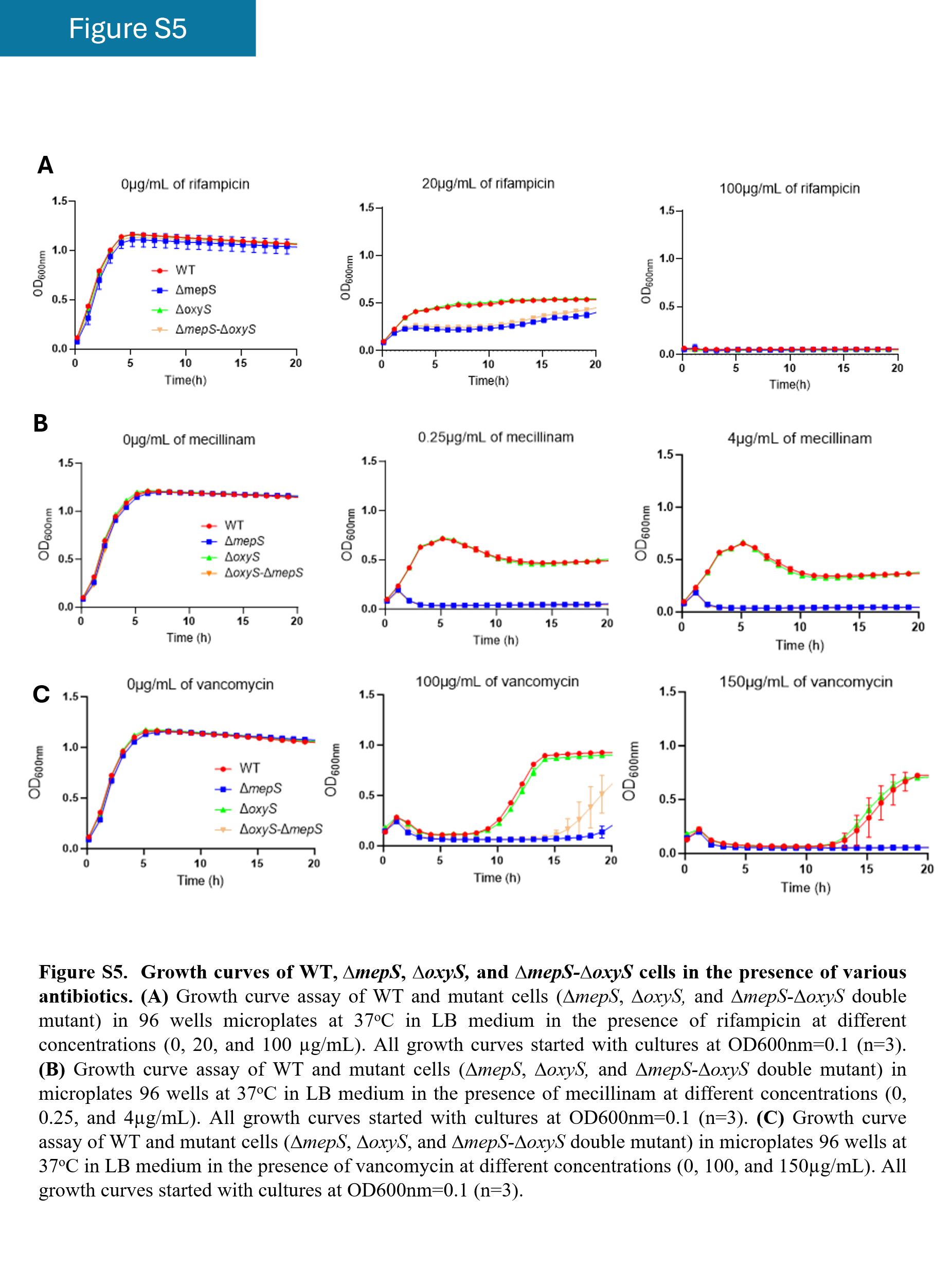

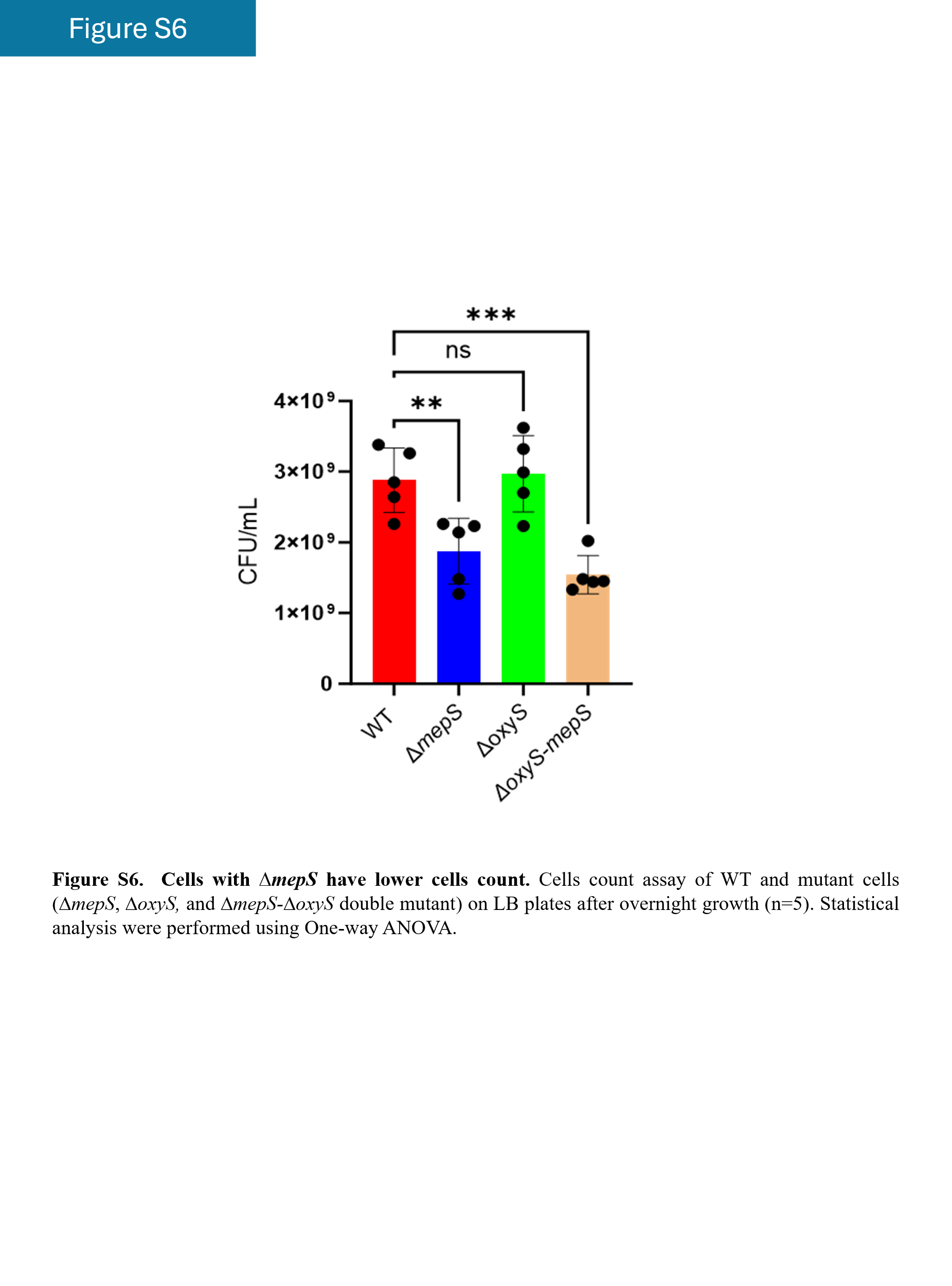

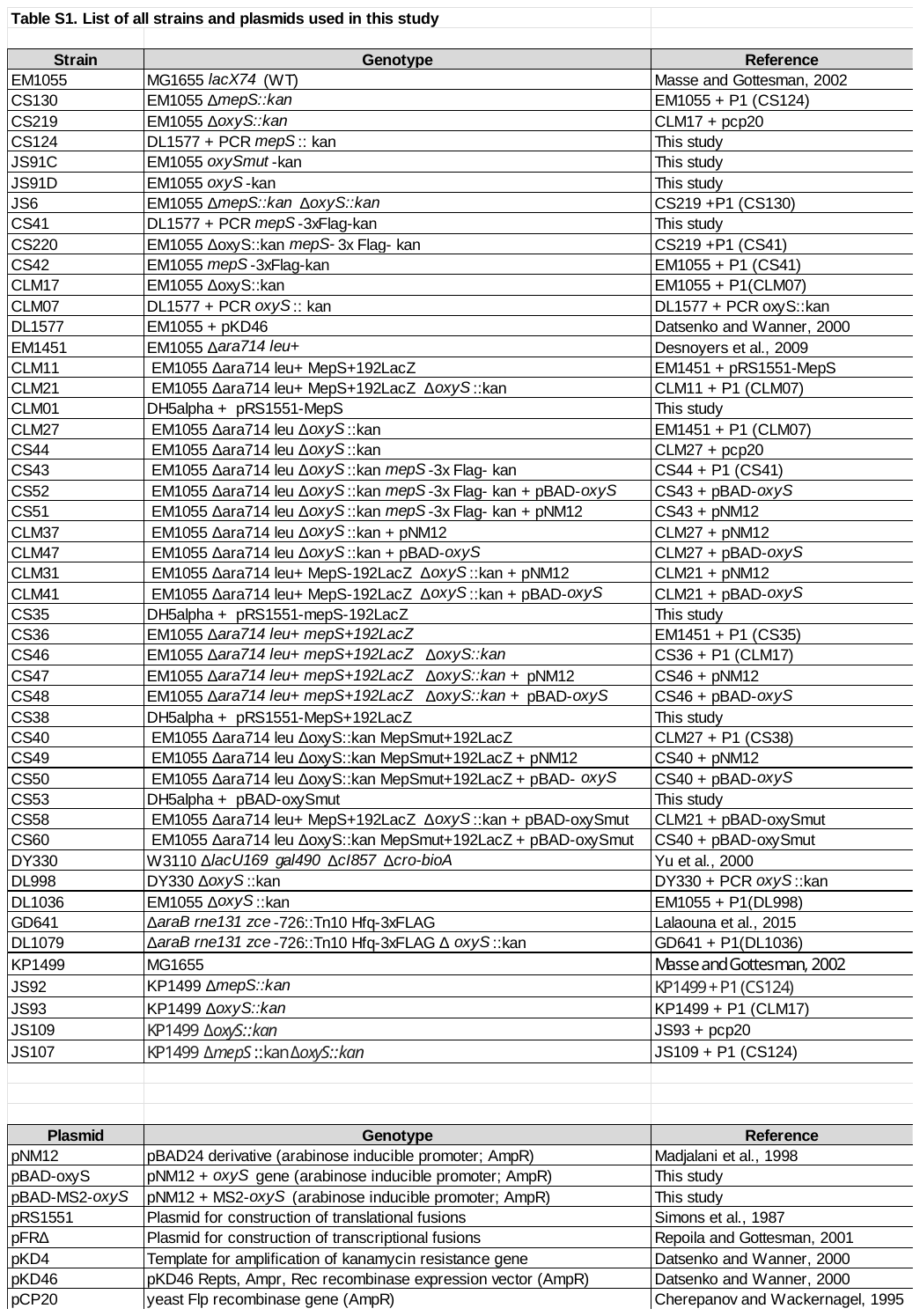

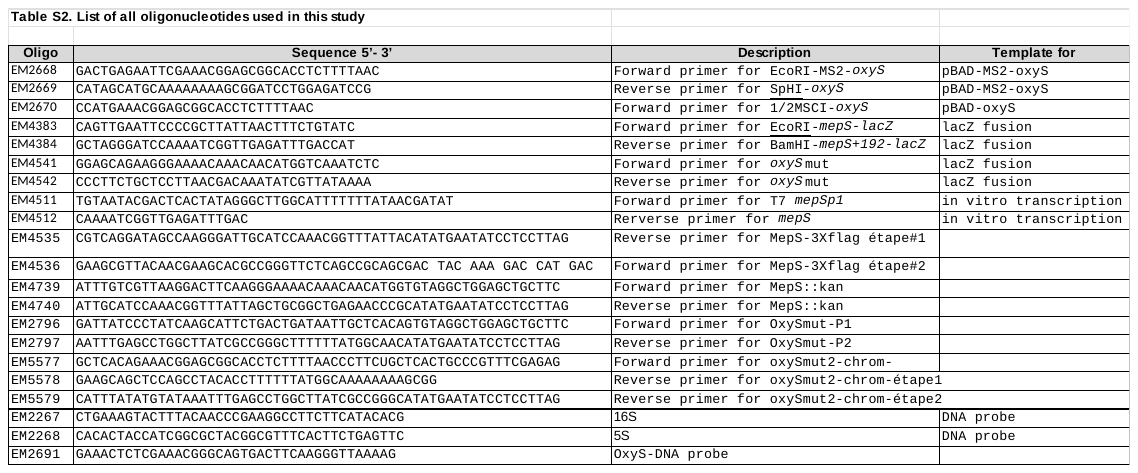
